# A method for the detection and enrichment of endogenous cereblon substrates

**DOI:** 10.1101/2025.03.24.645063

**Authors:** Hannah C. Lloyd, Yuli Li, N. Connor Payne, Zhenguang Zhao, Wenqing Xu, Alena Kroupova, David Zollman, Tengfang Long, Farah Kabir, Mei Chen, Rebecca Freeman, Ethan Yang Feng, Sarah Xi, Ya-Chieh Hsu, Alessio Ciulli, Ralph Mazitschek, Christina M. Woo

## Abstract

C-Terminal cyclic imides are posttranslational modifications on proteins that are recognized and removed by the E3 ligase substrate adapter cereblon (CRBN). Despite the observation of these modifications across the proteome by mass spectrometry-based proteomics, an orthogonal and generalizable method to visualize the C-terminal cyclic imide would enhance detection, sensitivity, and throughput of endogenous CRBN substrate characterization. Here we develop an antibody-like reagent, termed “cerebody,” for visualizing and enriching C-terminal cyclic imide-modified proteins. We describe the engineering of CRBN derivatives to produce cerebody and use it to identify CRBN substrates by Western blot and enrichment from whole cell and tissue lysates. CRBN substrates identified by cerebody enrichment are mapped, validated, and further characterized for dependence on the C-terminal cyclic imide modification. These methods will accelerate the characterization of endogenous CRBN substrates and their regulation.

## Introduction

Cereblon (CRBN) is a substrate recognition adapter of the CRL4^CRBN^ E3 ubiquitin ligase complex that mediates protein selection for ubiquitination and degradation. CRBN was first described in association with neurological development in humans^1^ and subsequently identified as a primary target of thalidomide, lenalidomide, and pomalidomide, ligands that bind to the conserved thalidomide-binding domain (TBD) of CRBN^2,3^ (**Figure 1a–c**). Induced substrate degradation partially underlies the observed therapeutic efficacy of thalidomide and its derivatives in multiple myeloma^4,5^ and del(5q) myelodysplastic syndrome,^6^ as well as off-target teratogenic effects.^7–9^ These findings have sparked immense efforts in the field of targeted protein degradation to induce a range of substrates to CRBN for therapeutic benefit.^10,11^ By contrast, the endogenous substrates mediated by CRBN and how they are affected by thalidomide and its derivatives remain relatively underexplored.

**Figure 1.**
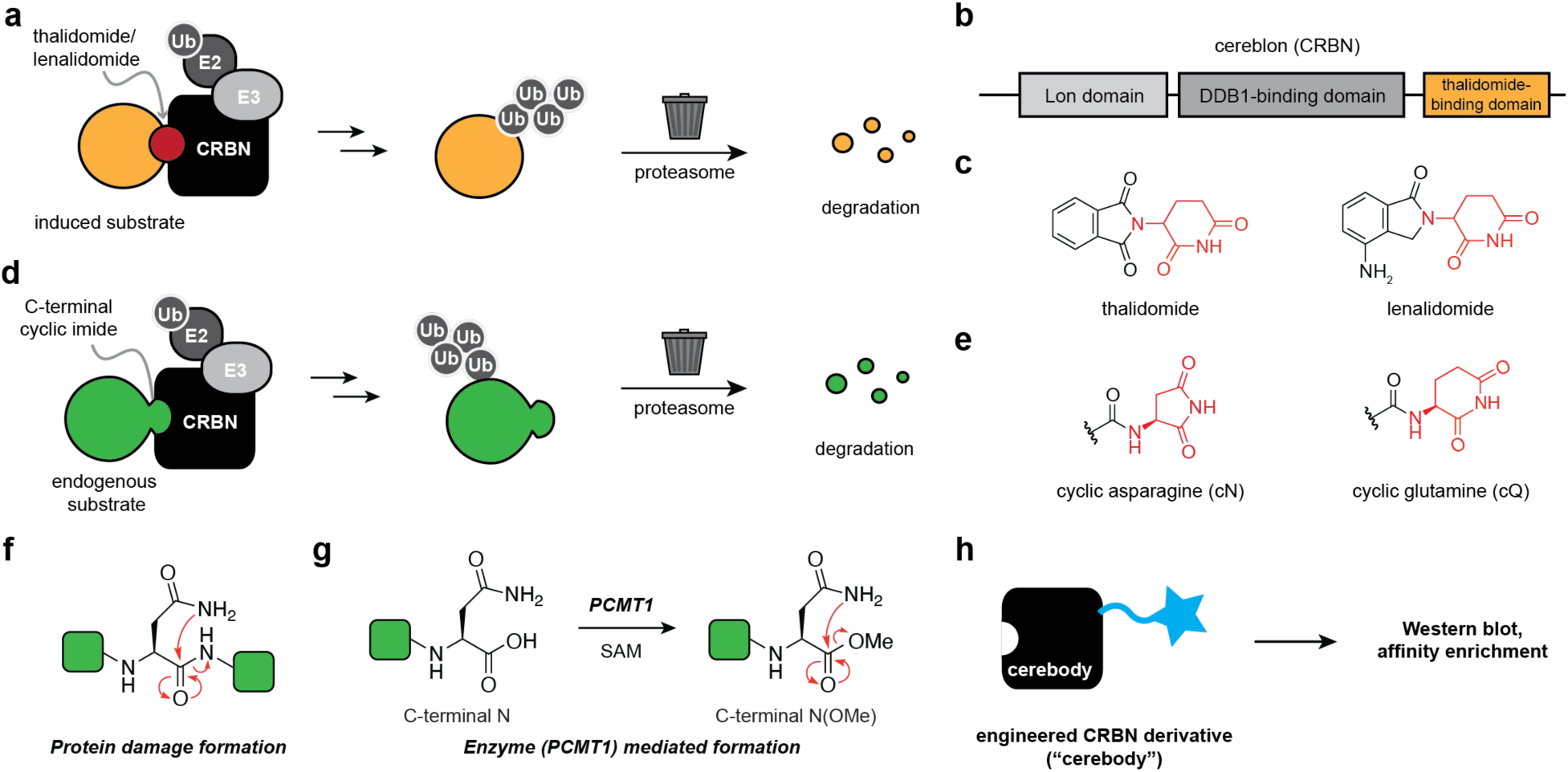
Development of a methodology to detect, image, and enrich C-terminal cyclic imides. **(a)** Thalidomide, lenalidomide, and derivatives induce substrates to CRBN for ubiquitination and subsequent degradation. **(b)** Linear schematic of domains in CRBN. **(c)** Structures of thalidomide and lenalidomide. **(d)** Endogenous substrates bearing C-terminal cyclic imides are recognized for ubiquitinylation by CRBN, leading to their subsequent degradation. **(e)** Structure of the C-terminal cyclic imide degron from cyclization of asparagine (cN) or glutamine (cQ). **(f–g)** C-Terminal cyclic imides can be formed by protein damage **(f)** or enzymatic **(g)** mechanisms. **(h)** Applications of an ideal “cerebody” for CRBN substrate recognition by Western blot or affinity enrichment.

Studies on the physiological function of CRBN have begun to reveal the molecular determinants for endogenous substrate recognition through the TBD. Our investigation of endogenous chemical motifs that thalidomide mimics led to the identification of the C-terminal cyclic imide modification that constitutes the minimal recognition motif, or degron, used by CRBN to recognize and remove substrates (**Figure 1b–e**).^12^ Coincident efforts provided a structural basis for C-terminal cyclic imide binding to the TBD.^13^ The C-terminal cyclic imide primarily arises from cyclization of asparagine, wherein the side chain amide attacks the carbonyl in the protein backbone, leading to generation of the C-terminal aspartimide (cN, **Figure 1f**). This modification can be formed as a result of intramolecular cleavage of the protein backbone during protein aging or stress, as occurs on glutathione synthetase (GSS),^14^ or via an enzymatic pathway for conversion of the C-terminal asparagine residue of proteins by protein carboxymethyltransferase (PCMT1) that go on to become substrates for CRBN (**Figure 1g**).^15^ The latter emerging pathway wherein PCMT1 acts on C-terminal asparagine to promote cyclic imide formation provides a molecular basis for prior studies on endogenous substrates that interface with the TBD, such as glutamine synthetase (GLUL).^15,16^ Thus, further investigation of the regulation of substrates by PCMT1 and CRBN through the C-terminal cyclic imide may illuminate deleterious effects on the brain and other phenotypes resulting from loss-of-function of PCMT1, CRBN, or their substrates, as well as their contribution to desirable or undesirable effects of thalidomide and its derivatives.

A generalizable method to visualize endogenous CRBN substrates that bear the C-terminal cyclic imide modification is critical to accelerate these investigations. However, methods for detecting C-terminal cyclic imides are currently confined to mass spectrometry analyses. Typically, the C-terminal cyclic imide can be detected by searching global proteomics datasets for semitryptic peptides with a dehydrated C-terminal asparagine or glutamine residue.^12,14^ However, these modifications may occur on peptides that are not ideal for trypsin digestion, the analyses must be controlled for mass spectrometry artifacts,^17^ and sensitivity is likely impacted by detection limits in the absence of an enrichment method. Furthermore, the C-terminal cyclic imide is prone to hydrolysis with a protein half-life of approximately 24 h,^12^ which can be lost during lengthy preparation protocols. Hydrolysis of the cyclic imide is further accelerated by basic conditions, such as those typically used in sample preparation for global proteomics experiments.^14^ By contrast, analyses of many protein modifications have been dramatically accelerated by modification-specific antibodies to detect, enrich, or image the modification in the proteome.^18^ We therefore sought to develop an antibody-like reagent for C-terminal cyclic imides to accelerate the study of this overlooked modification on endogenous substrates of CRBN.

In defining criteria for our antibody-like reagent, we considered the features of successful antibodies: good binding affinity to our epitope (K_D_ under 1 μM), stability in Western blot and enrichment experimental conditions, a slow off-rate with the C-terminal cyclic imide to maintain interactions through washing steps, the feasible ability to obtain sufficient quantities, and compatibility with other commercially available reagents. Furthermore, a C-terminal cyclic imide detection reagent should be specific to the C-terminal cyclic imide while remaining agnostic to the preceding amino acids.^19^ We considered two approaches to develop such a tool: traditional antibody generation by *in vivo* immunization and protein engineering of a known C-terminal cyclic imide binder. In considering the first approach, wherein the typical procedure for development of an antibody involves immunizing an animal with the desired target over the course of multiple weeks and then purifying an antibody from the plasma,^20^ we were uncertain about the expected success given the lability of the C-terminal cyclic imide to hydrolysis combined with the challenge of producing a reagent that recognizes C-terminal cyclic imides in a manner agnostic to the preceding sequence.^21^ Therefore we pursued the latter protein engineering approach to develop an antibody-like reagent from the C-terminal cyclic imide binder CRBN and demonstrate its use in Western blot detection and affinity enrichment of C-terminal cyclic imide bearing proteins from various cell and tissue lysates (**Figure 1h**).

## Results

### Development of cerebody by protein engineering

To begin engineering a known C-terminal cyclic imide binder into a suitable tool for detecting CRBN substrates, we first considered a series of CRBN constructs for their ability to be turned into a method for detection of CRBN substrates (**Figure 2a**). CRBN alone is unstable but is readily expressed in complex with DDB1, which has previously been used for affinity enrichment procedures to identify induced substrates of thalidomide derivatives.^4^ The CRBN/DDB1 complex has a good basal affinity to C-terminal cyclic imides, with a K_D, CRBN/DDB1_ = 96 +/– 21 nM to thalidomide-FITC (thal-FITC) by time-resolved Forster resonance energy transfer (TR-FRET) (**Figure 2b**, **Extended Data Figure 1a**), but is recombinantly expressed in insect cells and is unstable to *E. coli* overexpression, making it relatively challenging to readily obtain in large quantity. The TBD alone has a lower basal affinity to C-terminal cyclic imides, with K_D, TBD_ = 253 +/– 38 nM to thal-FITC, but is able to be recombinantly produced in *E. coli* and is the endogenous minimal binding motif to CRBN’s substrates through the cyclic imide degron.^22^ A third construct, CRBNmidi,^23^ was developed to recapitulate the biochemical properties of CRBN while avoiding the expression system liabilities of the CRBN/DDB1 complex. CRBNmidi has a strong basal affinity to thal-FITC, with K_D, CRBNmidi_ = 76 +/– 8 nM, and is readily overexpressed in *E. coli*, but includes 12 point mutations compared to human CRBN, as well as truncations of the DDB1 binding domain and N and C termini, although all of these changes are distal to the thalidomide binding pocket (**Figure 2a–b**).^23^ We examined the basal affinity of CRBN ligands thal-FITC and GFP-GLUL-cN, a representative endogenous substrate generated by reaction of GFP-GLUL with PCMT1, against these three CRBN constructs by TR-FRET to find similar trends (**Extended Data Figure 1b**, **Figure 2c**).^15^ We thus investigated each of these CRBN constructs as starting points for protein engineering campaigns to develop a robust tool for detecting C-terminal cyclic imides, using thal-FITC as a reasonable proxy for broad engagement of C-terminal cyclic imides.

**Figure 2.**
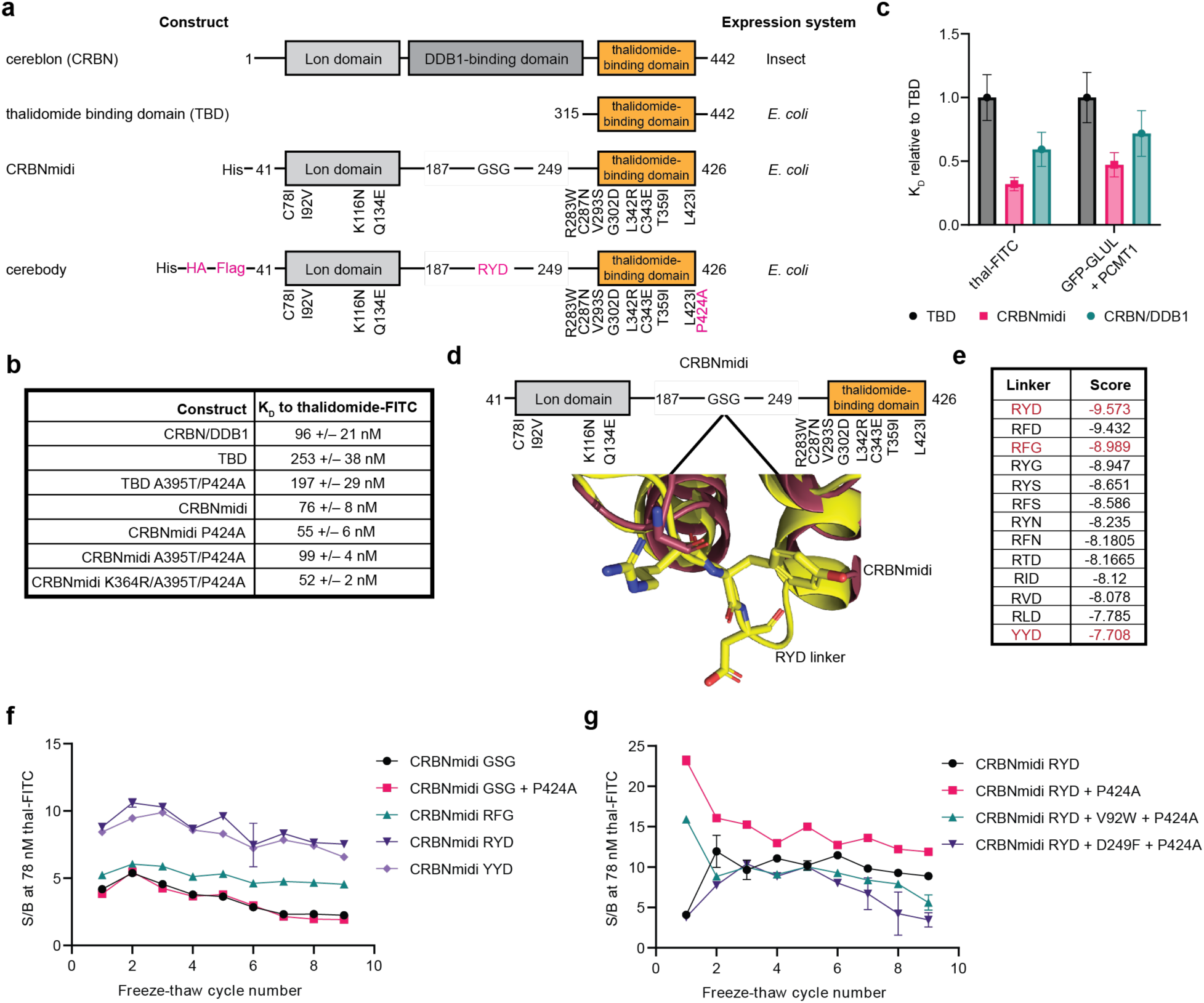
Development of cerebody, an antibody-like reagent for detecting C-terminal cyclic imides. **(a)** Full length human CRBN, the thalidomide binding domain, CRBNmidi, and cerebody. Changes from CRBNmidi to cerebody highlighted in pink. **(b)** Dissociation constant (K_D_) of indicated constructs to dose response of thal-FITC by TR-FRET. **(c)** Relative dissociation constants of TBD, CRBN/DDB1, and CRBNmidi to thal-FITC and GFP-GLUL treated with PCMT1. **(d)** Linker region of CRBNmidi with visualization of the RYD linker (yellow) overlaid on the crystal structure of CRBNmidi bound to lenalidomide (maroon, PDB 8RQA). **(e)** Sampling of lowest predicted linker energies by computational modeling. Experimentally examined linkers are highlighted in red. **(f)** Calculated signal to background ratios of CRBNmidi with noted linker residues against thal-FITC (78 nM) across 9 freeze-thaw cycles. **(g)** Calculated signal to background ratios of CRBNmidi with RYD linker containing the indicated mutation against thal-FITC (78 nM) across 9 freeze-thaw cycles.

We first investigated full-length CRBN/DDB1 for compatibility as a primary antibody with a HRP-conjugated α-His antibody as a secondary for Western blot against an engineered green fluorescent protein (GFP) that displays the C-terminal cyclic imide modification. This protein is generated by the sortase-catalyzed reaction of GFP-LPETG with a chemically-defined synthetic peptide (**Extended Data Figure 1c**).^24,25^ We quickly found that although the CRBN/DDB1 complex is readily used to measure substrate binding in solution using assays like TR-FRET,^19^ it is not compatible with Western blot (**Extended Data Figure 1d**). Furthermore, the His-tag is not well suited for selective detection of CRBN/DDB1 in complex cell lysates due to many other cross-reactive proteins, but the insect cell expression of CRBN/DDB1 placed limitations on our exploration of additional tags and the overall quantity of recombinant protein that could be obtained for broad use of the method. These limitations with CRBN/DDB1 led us to investigate other constructs.

We next investigated the TBD of CRBN, which has a relatively weak but measurable K_D_ to C-terminal cyclic imides and can be overexpressed in *E. coli* in scalable amounts (around 2 mg/L). Indeed, comparison of the binding of a thal–FITC probe to either CRBN/DDB1 or the TBD showed an approximate 2.5-fold difference between the constructs, which is within the range of binding affinity that could be recovered by directed evolution.^26^ We therefore considered the TBD as a good candidate for an affinity maturation campaign performed by random mutagenesis with a phage display library. We created a library of TBD derivatives with random mutations at a frequency of 9 mutations/kb fused to the N-terminus of pIII of filamentous phage.^27^ Phage were selected by in solution affinity to GFP-LcN, GFP-LcQ, GFP-FcN, GFP-FcQ, or GFP-PcQ (**Extended Data Figure 1e**). Mutations appearing at a variant frequency above 10% in the third and fourth rounds of selection were considered for validation, with preference given to variants that appeared in multiple selection conditions (**Supplemental Tables 1–3**). We individually overexpressed and purified hits selected for validation. Each TBD mutant was labeled with CoraFluor-1^28^ and examined for binding affinity to thal-FITC by TR-FRET. While calculated K_D_ values varied across experimental preparations, trends of constructs’ relative binding affinities remained largely consistent across experiments. For the first round, we selected 11 hits to characterize and found that the P424A mutation improved binding affinity to thal-FITC (**Supplemental Table 4**). A second round of phage display on a library based on TBD P424A yielded nine double mutants that were examined, with the double mutant A395T/P424A showing the strongest improvement in binding affinity to thal-FITC (**Supplemental Table 4**). A third round of phage display with a library based on TBD A395T/P424A did not produce any mutations at a variant frequency above 10%, and examination of six triple mutant TBD derivatives with a variant frequency between 5 and 10% did not yield a derivative that further enhanced the binding affinity (**Supplemental Table 4**). These efforts thus resulted in the identification of the double mutant TBD A395T/P424A with an approximately 2-fold improvement in binding affinity, which is at the upper edge of the range that is usable for enrichment.^26^

We next turned to investigation of the CRBNmidi scaffold^23^ given its retention of a binding affinity to CRBN ligands comparable to the CRBN/DDB1 complex due to the inclusion of the Lon domain,^29^ while being readily expressed in high yield in *E. coli* (approximately 1.5 mg/L,^23^ **Figure 2a**). We first investigated the translatability of mutations identified to enhance binding of the TBD and found that the P424A mutation slightly improved the dissociation constant to thal-FITC (**Figure 2b**). We installed Flag and HA tags at the N-terminus of CRBNmidi for compatibility with additional commercial detection reagents, and adapted the TR-FRET experiments to use a terbium labeled α-His antibody as previously reported,^28^ as this would be more representative of applications with commercial products targeting epitopes at the N-terminus of the construct. These modifications to CRBNmidi resulted in a protein compatible with a range of commercial reagents and an improved affinity to thal-FITC compared to that of CRBN/DDB1.

While CRBNmidi P424A produced a strong binder to thal-FITC, we observed variability in our downstream experiments, which we attributed to instability of the CRBNmidi construct. CRBNmidi and CRBNmidi P424A had a propensity for precipitation after more than one freeze-thaw cycle as measured by a decrease of signal and maximum signal-to-background ratio by TR-FRET, although the soluble fraction of the protein remained suitable for TR-FRET binding assays (**Extended Data Figure 2a–c**). We hypothesized that the flexible movement between the open and closed conformations of CRBNmidi^23^ could be a factor in its apparent instability as a reagent and therefore explored replacement of the flexible GSG linker connecting the Lon domain and TBD of CRBNmidi as a strategy to stabilize the protein (**Figure 2d**). We performed a virtual saturation mutagenesis screen of all tripeptide combinations of the 20 proteogenic amino acids that could be substituted at the position of the GSG linker (8000 combinations) to stabilize the closed conformation. Larger residues at the first and second linker positions scored well, while aspartate and glycine scored well at the third position. We selected RYD, RFG, and YYD for closer examination as substitutes for the GSG linker (**Figure 2e**). While the dissociation constant remained consistent across these constructs (**Extended Data Figure 2a**), we found the RYD linker had the best and most stable sensitivity to thal-FITC over nine freeze-thaw cycles (**Figure 2f**, **Extended Data Figure 2**). Installation of the P424A mutation further stabilized the resulting CRBNmidi-RYD-P424A construct to freeze-thaw cycles (**Figure 2g**). Parallel efforts with phage display selection of a randomly mutagenized library based on CRBNmidi-GSG-P424A against GFP-PFQYKcN, an engineered GFP that displays the C-terminal hexapeptide of GLUL, did not yield a stronger construct (**Extended Data Figure 3**). Additional computational modeling to identify potentially advantageous mutations at the interface of the Lon domain and the TBD was not productive (**Extended Data Figure 3**). We therefore completed the engineering of CRBN constructs to identify a more stable Flag and HA-tagged CRBNmidi-RYD-P424A derivative that has strong binding capacity for thal-FITC, which is henceforth referred to as “cerebody”.

### Detection of CRBN substrates by Western blot

We next examined cerebody’s utility as a primary antibody in Western blotting experiments (**Figure 3a**). We assessed its ability to detect C-terminal cyclic imides on proteins previously demonstrated to be degraded in a CRBN-dependent manner, including GLUL-cN generated by the reaction of recombinant GLUL with PCMT1^15^ or C-terminal cyclic imides installed on GFP^12^ or GSS^14^ by eSrtA-catalyzed reactions (**Extended Data Figure 4a**, **Figure 3b-c**). Using these protein standards, we examined detection efficiency against serial dilutions of GFP-PFQYKcN, GSS-cN, and GLUL-cN when spiked into HEK293T lysates (**Extended Data Figure 4b**). We determined that cerebody is best used at a working concentration of 5 μM with 5% nonfat dried milk in TBST as the blocking buffer with a HRP-conjugated α-HA antibody at 1:1000 dilution in blocking buffer as the secondary antibody (**Extended Data Figure 4b**). These conditions show a limit of detection between 3.2 x 10^−13^ and 6.4 x 10^−14^ mol of protein, comparable to the approximately 10^−12^ mol limit of detection of antibodies developed for other PTMs^30^ (**Figure 3d**).

**Figure 3.**
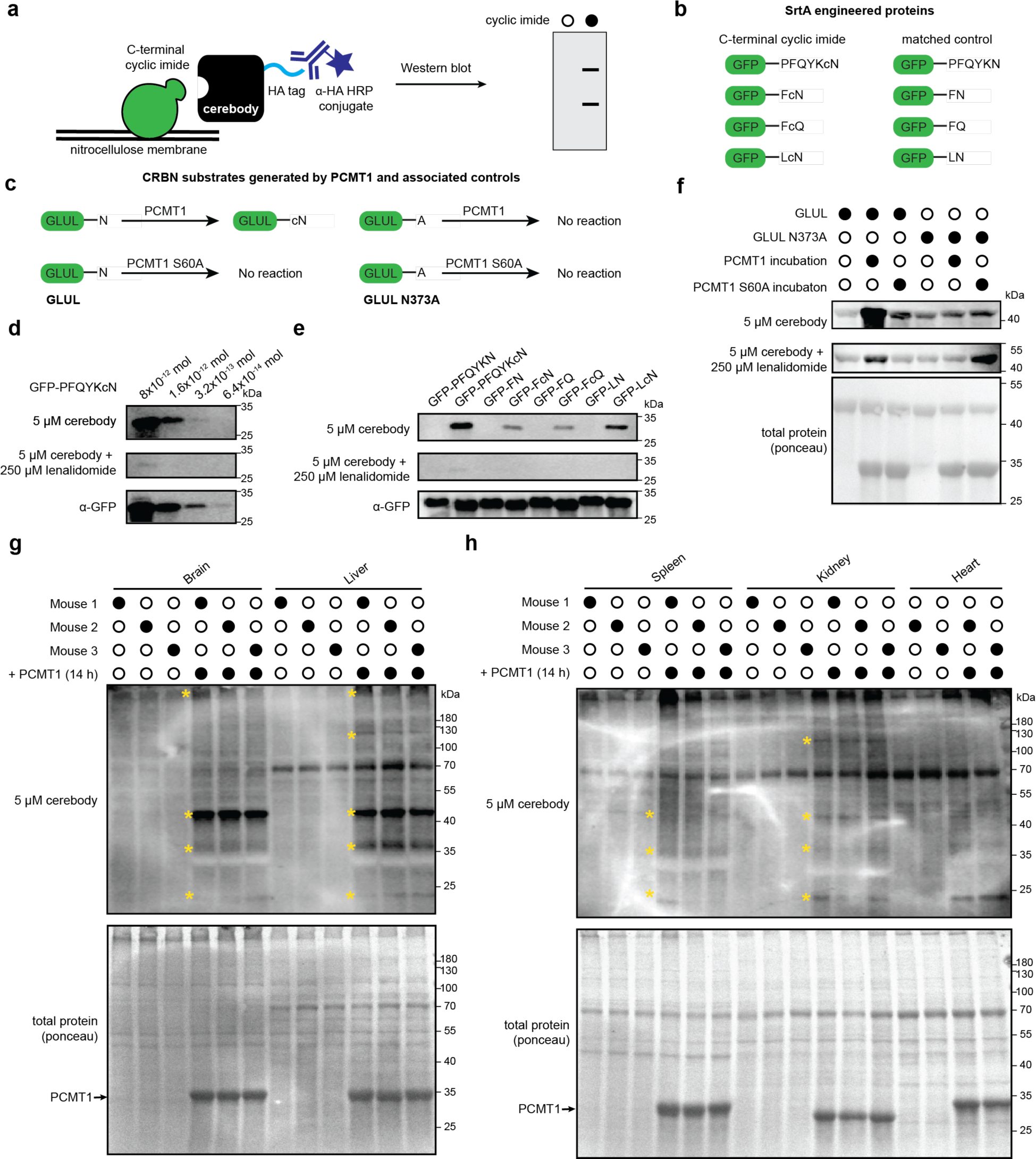
Western blotting for C-terminal cyclic imides using cerebody reagent. **(a)** Illustration of Western blot protocol with cerebody. **(b)** Engineered GFP displaying C-terminal cyclic imide or matched control generated by SrtA. **(c)** Schematic of CRBN substrate GLUL-cN generated by PCMT1 with associated controls. **(d)** Limit of detection of cerebody measured by serial dilution of GFP-PFQYKcN. **(e)** Cerebody detection of the indicated engineered GFP construct at 8 x 10^−12^ mol. **(f)** Cerebody blot of recombinant GLUL or GLUL N373A following treatment with PCMT1 or inactive PCMT1 S60A control. **(g)** Cerebody blot of mouse brain and liver lysates with or without PCMT1 incubation (14 h). Bands appearing following PCMT1 incubation highlighted with yellow asterisks. **(h)** Cerebody blot of mouse spleen, kidney, and cardiac tissue lysates with or without PCMT1 incubation. Bands appearing following PCMT1 incubation highlighted with yellow asterisks.

In each instance, the detection of the engineered protein was dependent on the presence of the C-terminal cyclic imide (**Figure 3e**). GLUL-cN had a much higher cerebody blotting signal compared to that of GLUL or GLUL-N373A that was ablated when 250 μM lenalidomide was present with cerebody during primary incubation (**Figure 3f**). Cerebody is therefore analogous to a primary antibody in Western blotting to visualize CRBN substrates bearing C-terminal cyclic imide PTMs in complex mixtures.

To examine the potential of cerebody Western blots to visualize C-terminal cyclic imides in whole cell lysates and in tissue samples, mouse tissue and HEK293T lysates were treated with exogenous PCMT1 for 14 h to promote formation of C-terminal cyclic imides on substrate proteins. Western blot analysis with cerebody revealed the presence of specific bands from different tissues that appeared in a PCMT1 dependent manner (**Figures 3g–h**). Several new bands, indicative of potential CRBN substrates, were particularly prominent in the mouse brain and liver when treated ex vivo with PCMT1, with additional bands observed across other examined tissues (highlighted by yellow asterisks, **Figure 3g–h**). In wild type HEK293T (HEK293T WT) or PCMT1 knockout HEK293T (HEK293T PCMT1 KO) cells, a band between 35 and 40 kDa is clearly observed following PCMT1 treatment, which is around the expected molecular weight for GLUL (**Extended Data Figure 4b**). Thus, cerebody enables visualization of specific bands, particularly after incubation with PCMT1, from cell culture and mouse tissue lysates.

### Enrichment of CRBN substrates from complex lysates

Given the efficient detection of established and candidate CRBN substrates by Western blot, we next explored the capability of cerebody to enrich CRBN substrates (**Figure 4a**). We first established the feasibility of cerebody affinity enrichment using HEK293T WT lysates spiked with GLUL-cN. GLUL-cN, but not GLUL, was selectively enriched by cerebody from these lysates (**Extended Data Figure 5a**).

**Figure 4.**
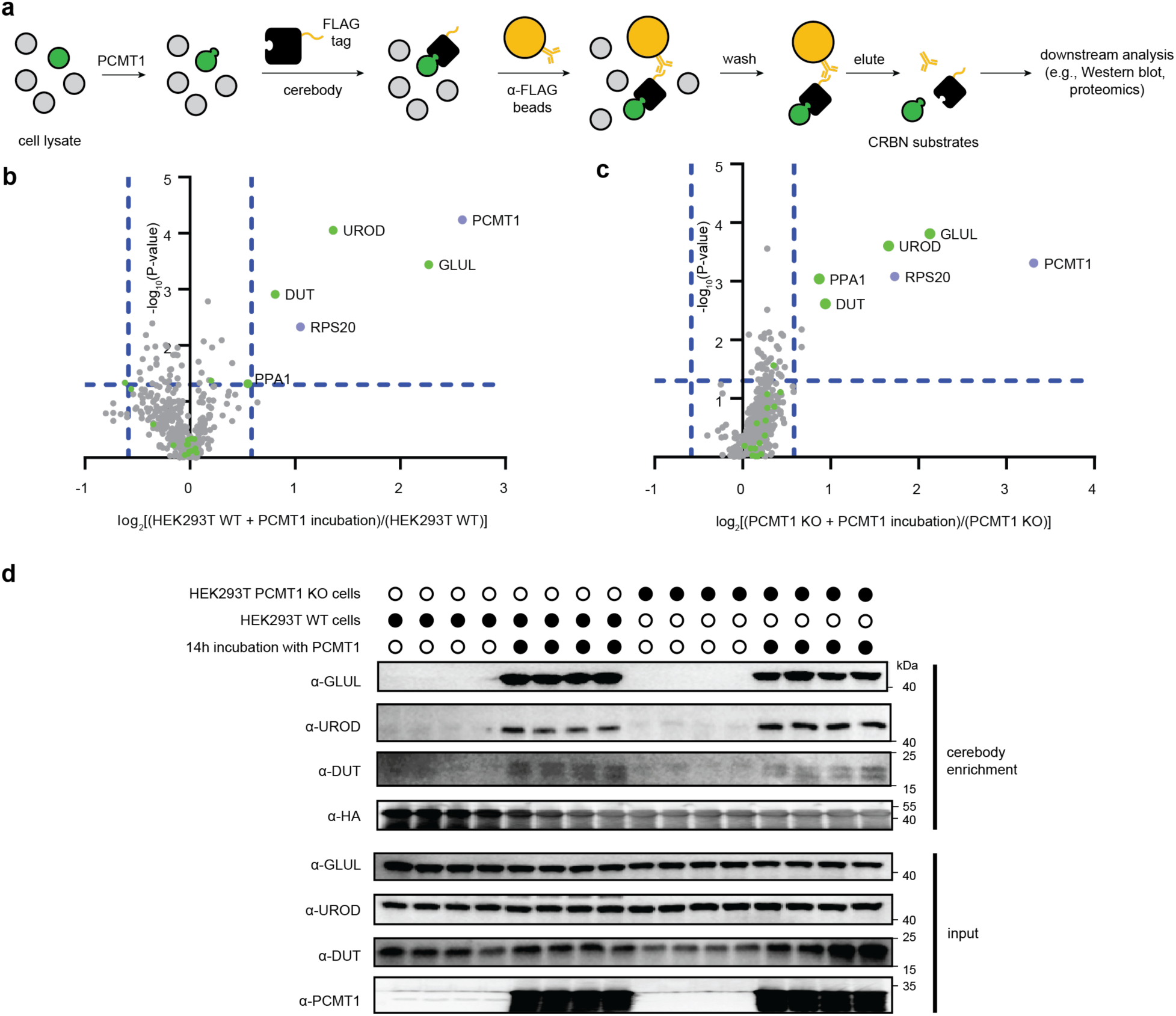
Affinity enrichment of candidate CRBN substrates with cerebody from HEK293T cell lysates. **(a)** Illustration of affinity enrichment procedure with cerebody. **(b)** Mass spectrometry analysis of HEK293T WT lysates with or without PCMT1 treatment. Green dots represent proteins with C-terminal asparagine. **(c)** Mass spectrometry analysis of HEK293T PCMT1 KO cells with or without PCMT1 treatment. Green dots represent proteins with C-terminal asparagine. **(d)** Western blot of proteins following affinity enrichment with cerebody. HEK293T WT or PCMT1 KO were treated with or without PCMT1 for 14 h.

To further illustrate the power of this method for discovery, we analyzed lysates from HEK293T WT or PCMT1 KO cells following exogenous PCMT1 incubation and cerebody affinity enrichment by mass spectrometry. Cell lysates were incubated with or without PCMT1 for 14 h prior to cerebody enrichment. Enriched lysates were trypsin digested and characterized by mass spectrometry. Datasets were assigned by a Proteome Discoverer search against the Human Swissprot dataset, and ratios calculated by comparing the proteins enriched from lysates with PCMT1 incubation to those enriched in the absence of PCMT1 incubation.

The two datasets showed remarkably similar cerebody enrichment profiles. Five proteins and PCMT1 were significantly enriched (fold-change ≥ 1.5, p-value ≤ 0.05) in both HEK293T WT and PCMT1 KO cells (**Figure 4b–c**). In both cell types, GLUL showed the highest fold change when comparing the PCMT1 treated lysates to those not treated with PCMT1. Inorganic pyrophosphatase (PPA1), which is also a substrate of PCMT1 and CRBN,^15^ was identified as significantly enriched in HEK293T PCMT1 KO cells, and was enriched in HEK293T WT lysates following PCMT1 treatment, although slightly under the 1.5-fold threshold (**Figure 4b–c**). Two other proteins with C-terminal asparagine, uroporphyrinogen decarboxylate (UROD), which is involved in heme biosynthesis,^31^ and mitochondrial deoxyuridine 5’-triphosphate nucleotidohydrolase (DUT), which prevents misincorporation of uridine into DNA,^32^ were identified as significantly enriched in both cell types in the PCMT1-treated condition. Gratifyingly, GLUL, UROD, and DUT enrichment were additionally visualized by Western blot following cerebody enrichment of PCMT1-treated lysates (**Figure 4d**). The fifth enriched protein in both cell types, small ribosomal subunit protein uS10 (RPS20), does not have a C-terminal asparagine and did not validate by Western blot (**Extended Data Figure 5b**). More broadly, proteins bearing a C-terminal asparagine appeared to be generally enriched in PCMT1-treated HEK293T PCMT1 KO lysates (highlighted in green, **Figure 4b-c**). These data demonstrate that cerebody can enrich and enable discovery of potential CRBN substrates that are promoted by PCMT1.

We next examined lysates of mouse tissues with and without *ex vivo* treatment with PCMT1. Glul was significantly enriched by cerebody across mouse brain, kidney, liver, spleen, and cardiac tissue lysates in a manner dependent on PCMT1 treatment by mass spectrometry and Western blot validation (**Figure 5**). Inorganic pyrophosphatase 2 (Ppa2) was significantly enriched in the brain, liver, spleen, and cardiac tissue lysates treated with PCMT1 (**Figures 5a–f; 5i–j**). Ppa2 is homologous to Ppa1, which was identified as a substrate in mouse tissues, including the spleen (**Figure 5c**), both of which terminate with an asparagine residue.^15^ Peroxiredoxin-4 (Prdx4), which has previously been identified as a CRBN interactor,^33^ was significantly enriched in the PCMT1-treated liver tissue (**Figure 5b**), and peroxiredoxin-2 (Prdx2) was significantly enriched in PCMT1-treated cardiac and kidney tissue lysates (**Figure 5g–h**). Both peroxiredoxins have a C-terminal asparagine. A third peroxiredoxin, peroxiredoxin-1 (Prdx1), that lacks a C-terminal asparagine, was identified as significantly enriched in PCMT1-treated cardiac tissue lysate. These peroxiredoxins protect against oxidative stress.^34^ An additional protein with a C-terminal asparagine, BAG family molecular chaperone regulator 2 (Bag2), was identified as enriched from PCMT1-treated cardiac tissue lysate. This protein has a conserved C-terminal BAG domain which interacts with the ATPase domain of Hsp70 to modulate its interactors.^35^ Enriched proteins that do not have C-terminal asparagine are more tissue specific, including band 3 anion transport protein (Slc4a1) from brain tissue lysate and homogentisate 1,2-dioxygenase (Hgd) identified in kidney tissue lysate. These data illustrate the generalizability of cerebody enrichment from a range of sample types that expedites discovery of potential substrates of PCMT1 and CRBN, thus providing new insight to candidate CRBN substrates that may be dependent on the C-terminal cyclic imide degron.

**Figure 5.**
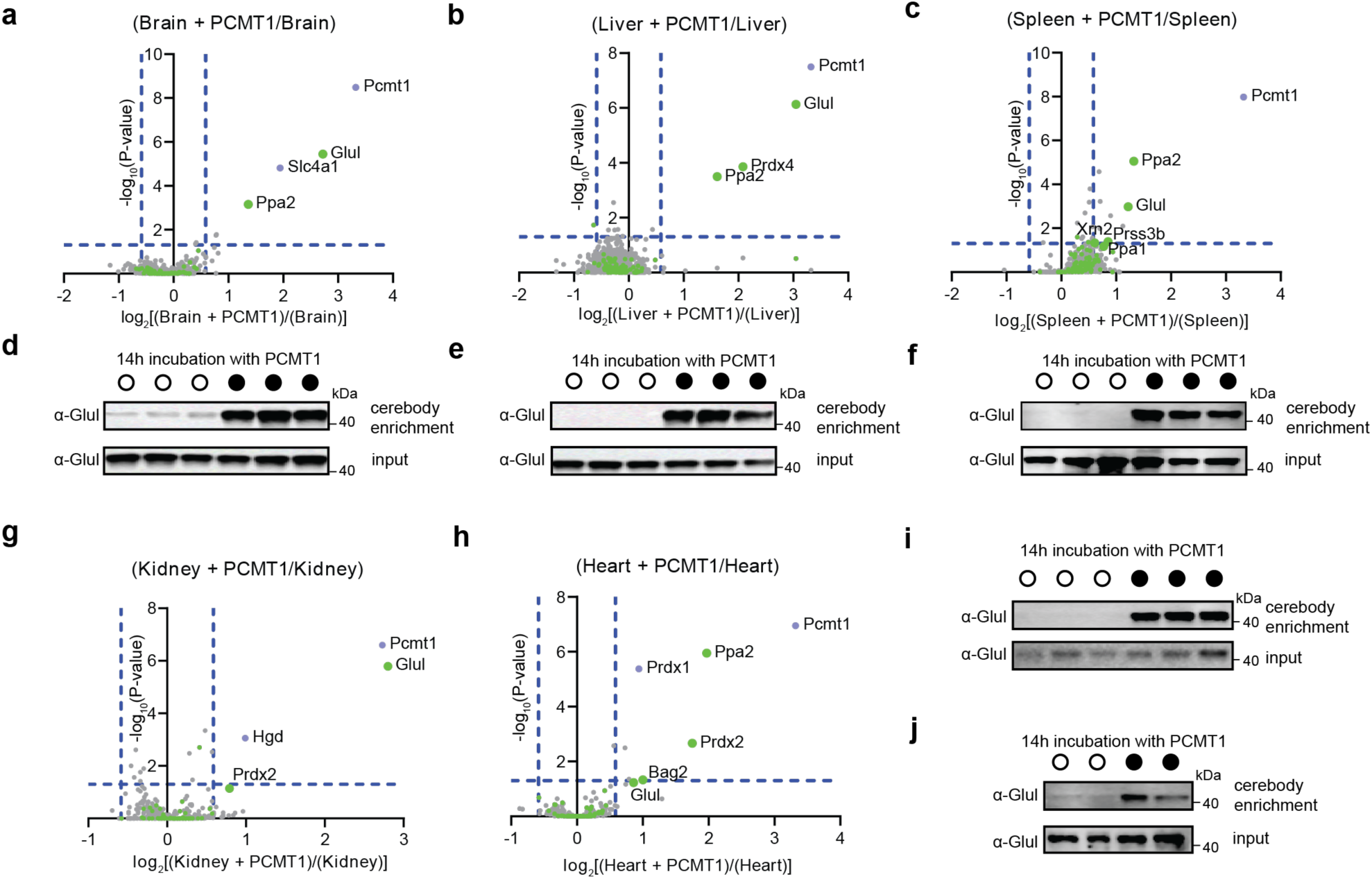
Analysis of mouse tissue lysates following PCMT1 treatment and cerebody enrichment. **(a–c)** Mass spectrometry analysis following cerebody enrichment of **(a)** brain, **(b)** liver, and **(c)** spleen tissue with or without PCMT1 incubation (14 h). Green dots represent proteins with C-terminal asparagine. **(d–f)** Western blot for GLUL following cerebody enrichment of **(d)** brain, **(e)** liver, and **(f)** spleen tissue. **(g–h)** Mass spectrometry analysis following cerebody enrichment of **(g)** kidney and **(h)** cardiac tissue with or without PCMT1 incubation (14 h). Green dots represent proteins with C-terminal asparagine. **(i–j)** Western blot for GLUL following cerebody enrichment of **(i)** kidney and **(i)** cardiac tissue.

### Identification of UROD as a PCMT1-dependent CRBN substrate

UROD is involved in the heme biosynthetic pathway, catalyzing the conversion of uroporphyrinogen to coproporphyrinogen (**Supplemental Figure 2**).^36^ Based on the enrichment and validation of UROD in PCMT1-treated conditions in both HEK293T WT and PCMT1 KO cells, we sought to further characterize UROD as a CRBN substrate via PCMT1-mediated C-terminal cyclic imide formation (**Figure 6a**). Recombinant UROD, but not UROD N367A, was recognized by cerebody on Western blot following PCMT1 treatment (**Figure 6b**). Treatment with catalytically inactive PCMT1 S60A did not promote signal by Western blot. The C-terminal cyclic imide on UROD was confirmed by mass spectrometry (**Figure 6c**). These data correspond to *in vitro* measurements by competitive TR-FRET against full length CRBN/DDB1 (**Figures 6d**). To assess dependence of UROD degradation on the C-terminal cyclic asparagine, GFP-UROD fusions with C-terminal asparagine or a C-terminal UROD N367A mutation were incubated *in vitro* with PCMT1 followed by electroporation to HEK293T WT or HEK293T CRBN knockout^12^ (CRBN KO) cells. GFP-UROD was degraded only in HEK293T WT cells, with rescue of degradation occurring with lenalidomide co-electroporation, where the N367A mutation and HEK293T CRBN KO cells did not demonstrate a significant difference in relative GFP levels after 6 h (**Figure 6e–f**). When the GFP-UROD fusions were electroporated into HEK293T WT cells without PCMT1 preincubation, a slight but significant decrease was observed between GFP-UROD with and without lenalidomide co-treatment, and between GFP-UROD and GFP-UROD N367A, while no significant difference was observed between the GFP-UROD N367A with and without lenalidomide co-electroporation with 42 h of incubation (**Figure 6g**). These data establish UROD is a substrate of PCMT1 and CRBN in vitro and following introduction to cells.

**Figure 6.**
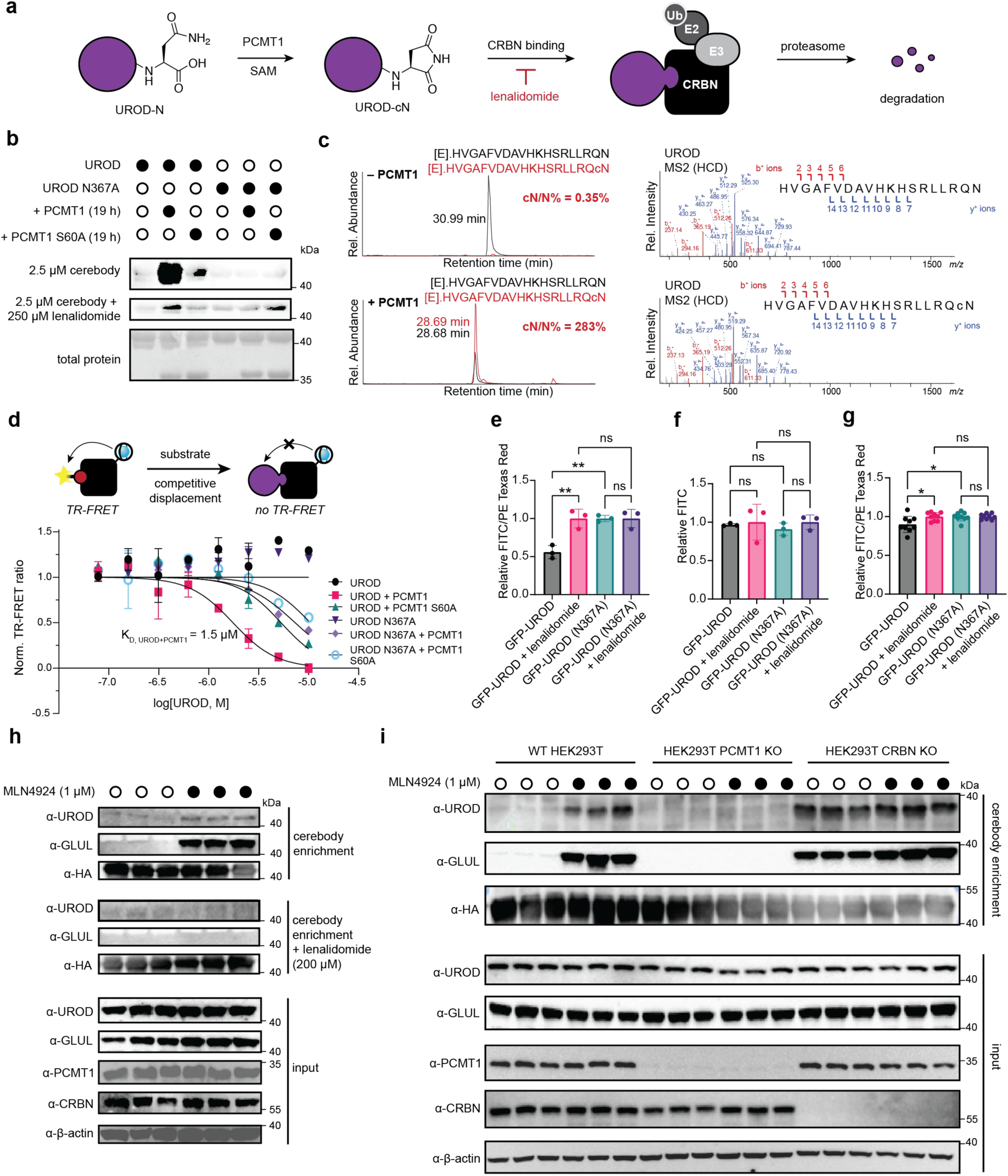
UROD is an endogenous substrate of CRBN. **(a)** Illustration of C-terminal cyclic imide formation on UROD leading to CRBN recognition and subsequent degradation. **(b)** Cerebody blot of UROD and UROD N367A incubated with or without PCMT1 or PCMT1 S60A for 19 h. **(c)** Selected ion monitoring and MS2 fragmentation spectra of the C-Terminal peptide of UROD treated with PCMT1 (17.5 h) and digested by Glu-C. **(d)** TR-FRET of UROD and UROD N367A following incubation with PCMT1 or PCMT1 S60A (17.5 h) by competitive displacement of thal-FITC against CRBN/DDB1. **(e–f)** Flow cytometry analysis of GFP-UROD incubated with PCMT1 (17.5 h) and electroporated into HEK293T WT **(e)** and CRBN KO **(f)** cells 6 h after electroporation. **(g)** Flow cytometry analysis of GFP-UROD following electroporation to HEK293T WT cells after 42 h. **(h)** Cerebody enrichment of HEK293T WT cells treated with MLN4924 or DMSO for 24 h prior to harvesting with DMSO or lenalidomide present during enrichment. **(i)** Blot of cerebody enrichment in HEK293T WT, PCMT1 KO, and CRBN KO cells with and without MLN4924 treatment (24 h) prior to harvesting.

We next used the cerebody method to confirm that endogenous UROD, like GLUL, is a substrate of CRBN. Both endogenous UROD and GLUL were affinity enriched by cerebody in HEK293T WT cells following treatment with the NEDD8-activating enzyme inhibitor MLN4924 (**Figure 6h**).

The enrichment was ablated by lenalidomide competition, indicating specificity of the enrichment depends on the TBD. Furthermore, examination of HEK293T WT, PCMT1 KO, and CRBN KO cells with and without MLN4924 treatment demonstrated that endogenous CRBN substrates GLUL and UROD build up when CRBN is lost or neddylation of Cullin Ring ligases is blocked, but not when PCMT1 is lost (**Figure 6i**). Together, these data illustrate the power of cerebody methods to discover and accelerate the biochemical characterization of endogenous CRBN substrates to rapidly establish substrates like UROD as regulated by PCMT1 and CRBN through the C-terminal cyclic imide degron.

## Discussion

Here we present the development of cerebody, a robust protein reagent to visualize and enrich CRBN substrates. CRBN and its substrates have been the subject of significant investigation due to their potential to be affected by CRBN-directed targeted protein degradation modalities^10,11^ and their important yet underexplored role in human biology.^1,37^ However, CRBN’s principle recognition motif of substrates through a protein modification, the C-terminal cyclic imide degron, has made these substrates challenging to visualize in a routine manner other than by global proteomics. By leveraging protein engineering approaches to derivatize CRBNmidi,^23^ we now report the development of cerebody as a reagent that renders routine the detection of CRBN substrates. We report the use of cerebody to detect, enrich, and discover proteins with C-terminal cyclic imides including engineered proteins, recombinant proteins, and endogenous CRBN substrates like GLUL and UROD. This method will accelerate the understanding of the native interactors of CRBN, the mechanisms through which they interact to shed light on CRBN’s physiological function, and the ways these pathways are impacted by CRBN ligands.

Use of CRBN as an enrichment modality^16,38,39^ along with a number of emerging technologies using proximity-based labeling have been employed to characterize CRBN interactors, primarily in the presence of small molecule ligands.^33,40–45^ Enrichment with CRBN/DDB1 can be a successful strategy to identify induced substrates and provide insights to endogenous substrates like GLUL,^16^ but use of this protein complex is relatively constrained by the expression system, not compatible with Western blot, and enrichments can be highly variable in our experience. Proximity-labeling strategies for proteins in proximity to CRBN or following protein ubiquitination in a CRBN-dependent manner are complementary approaches that are revealing profiles which include proteins like GLUL and PPA1 that are likewise observed in our cerebody enrichments.^40,42–44^ However, despite the power of proximity-based labeling methods, they require engineering of the cell for functionality. By contrast, cerebody detection and enrichment methods offer a generalizable and complementary approach to detect and enrich CRBN substrates in cell and tissue lysates without cellular engineering. These collective advances are strengthening connection between the C-terminal cyclic imide degron and endogenous substrates of CRBN.

As demonstration of the ease with which cerebody facilitates endogenous CRBN substrate characterization, our examination of cell and tissue lysate samples by cerebody enrichment led to the identification of both previously characterized substrates for PCMT1-mediated C-terminal cyclization, GLUL and PPA1,^15^ and of the newly characterized substrates DUT and UROD. We further biochemically characterize UROD as an endogenous substrate of PCMT1-mediated C-terminal cyclization and demonstrate the cellular relevance of this mechanism for CRBN-dependent degradation of C-terminally cyclized UROD. Cerebody enrichment has therefore strengthened the characterization of the role of C-terminal cyclic imides in these substrates’ interactions with CRBN and opens opportunities for exploring substrates of CRBN and their biological implications, thus providing exciting new directions in the investigation of the endogenous biology of both PCMT1 and CRBN.

## Methods

### Protein labeling for TR-FRET measurements

α-His_6_ antibody (Abcam 18184), His-CRBN/DDB1, TBD constructs, and CRBNmidi constructs were labeled with CoraFluor-1-Pfp ester as previously described.^28,46^ The following extinction coefficients were used to calculate protein concentration and degree-of-labeling (DOL): anti-His_6_ antibody *E*_280_ = 210,000 M^-1^cm^-1^, CRBN-DDB1 *E*_280_ = 167,000 M^-1^cm^-1^, TBD constructs *E*_280_ = 28,000 M^-1^cm^-1^, CRBNmidi constructs *E*_280_ = 55,350 M^-1^cm^-1^, CoraFluor-1-Pfp *E*_340_ = 28,000 M^-1^cm^-1^. Protein conjugates were stored at -80°C.

### TR-FRET measurements

Experiments were performed in Proxiplate-384 Plus (VWR PERK6008280) in 10 μL assay volume. TR-FRET measurements were acquired on a on a SpectraMax iD5 plate reader with SoftMax Pro software version 7.1.2, with the following settings: 350 nm excitation, 616 nm (Tb), and 520 nm (FITC) emission, 110 flashes per read, 0.05 ms excitation time, 0.05 ms delay and 0.2 ms integration or on a Tecan SPARK plate reader with SPARKCONTROL software version V2.1 (Tecan Group Ltd.), with the following settings: 340/50 nm excitation, 490/10 nm (Tb), and 520/10 nm (FITC, GFP) emission, 100 ms delay, 400 ms integration. The 490/10 nm and 520/10 nm emission channels were acquired with a 50% mirror and a dichroic 510 mirror, respectively, using independently optimized detector gain settings. The TR-FRET ratio was taken as the 520/490 nm or 520/616 nm intensity ratio on a per-well basis.

### TR-FRET affinity measurements

For ligand association assays, 50 nM CoraFluor-1-labeled TBD construct or CoraFluor-1-labeled CRBN/DDB1 complex, 25 nM CoraFluor-1-labeled CRBNmidi construct, or 10 nM unlabeled CRBNmidi construct and 5 nM CoraFluor-1-labeled α-His_6_ antibody was added to the assay buffer (25 mM HEPES, 150 mM NaCl, 0.5 mg/mL BSA, 0.005% TWEEN 20, pH 7.5) and homogenized. Freeze-thaw cycling was performed using unlabeled CRBNmidi that was placed at -80oC to solidify between cycles, and a fresh aliquot of CoraFluor-1-labeled α-His_6_ antibody from the same preparation was used for each cycle. Thal-FITC in DMSO or GFP proteins with 0.3% TWEEN-20 were added in serial dilution (1:2 titration, 9-point, c_max_ = 2.5 µM for thal-FITC, 1 µM for GFPs) using an HP D300 digital dispenser. The same serial dilution of each protein sample was performed twice, once without any additive as the sample well and again with 100 µM 5-NH_2_-lenalidomide added as the background well to compete off CRBN-specific association. All experiments were performed in technical triplicates. The corrected TR-FRET ratio was determined by subtracting the ratio of background well from that of the sample well containing the same concentration of GFP sample. Data was fitted to a one-site specific binding model in Prism 10 to derive the K_D_ values. Relative K_D_ values were calculated by dividing the calculated K_D_ by the calculated K_D_ of the relative construct as measured in the same experiment.

### TR-FRET displacement assay

For ligand displacement assays, 20 nM CoraFluor-1-labeled CRBN/DDB1 complex and 200 nM Thal-FITC were mixed in the assay buffer (25 mM HEPES, 150 mM NaCl, 0.5 mg/mL BSA, 0.005% TWEEN 20, pH 7.5) and 5 μL added to each well. UROD reaction mixtures with TWEEN-20 added to 0.3% (v/v) were added in serial dilutions (1:3 titration) to each well and the well volume brought to 10 μL using an HP D300 digital dispenser. All experiments were performed in technical triplicate. Signal was determined after 2 h. Background signal (bottom) was determined from wells containing the maximum concentration of UROD incubated with PCMT1 to represent complete displacement. The assay ceiling (top) was defined via a no-ligand control. Data were background-subtracted, normalized, and fitted to a 4-parameter dose-response model [log(inhibitor) vs. response – Variable slope (four parameters)] in Prism 10, with constraints of Top = 1 and Bottom = 0 to derive the ligand IC_50_ values. Ligand K_D_ values have been calculated using Cheng-Prusoff principles, outlined in the equation below:

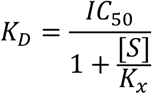

Where IC_50_ is the measured IC_50_ value, [S] is the concentration of fluorescent tracer, and K_X_ is the K_D_ of the fluorescent tracer without adjustment.

### Affinity maturation of TBD

Random mutagenesis was performed on target coding region using flanking primers 99 and 100 with SfiI restriction sites inserted using Agilent GeneMorph II kit (Agilent Technologies 200550). The randomly mutagenized insert and pCOMB3xSS vector were digested by SfiI (New England Biolabs R0123S) and the backbone gel extracted using the Monarch gel extraction kit (New England Biolabs T1020S). The backbone and insert were ligated at a 1:3 ratio using T4 DNA ligase (New England Biolabs M202T) at room temperature for 2 h. The ligation mix was transformed in 5-alpha (New England Biolabs C2987H) and random mutagenesis was verified by Sanger sequencing of the insert region of individual colonies using primers 101 and 102. Verified libraries were electroporated into XL1-Blue cells (Agilent 200228) at 1700V using an Eppendorf electroporator. Cells were allowed to grow for 30 minutes, at which point 30 μL helper phage (M13KO7 New England Biolabs N0315) were added and the cells allowed to grow for 1 hour. Additional media, kanamycin (50 μg/mL), and carbenicillin (100 μg/mL) were added and the cells allowed to grow overnight. Cells were pelleted at 4,000xg for 15 minutes at 4°C. Spent media was combined 5:1 with 2.5 M NaCl, 20% (w/v) PEG-8000 and incubated on ice for 20 minutes. Phage were pelleted at 12,000xg, 4°C, 30 minutes, and resuspended in PBS pH 7.4, 0.05% (v/v) Tween-20, 0.05% (w/v) bovine serum albimum. Resuspended phage was added to GFP-trap coupled to magnetic agarose (Bulldog Bio GTA020) preincubated for 1 hour with GFP previously reacted by sortase.^3^ Phage were incubated with end-over-end motion at 4°C for 1-2 h. Phage were washed as described in supplemental table and eluted by incubation with 0.1 M hydrochloric acid for 10 minutes at room temperature followed by neutralization with an equivalent volume of 1 M Tris pH 11 or by incubation with trypsin at 37°C for 1.75-3 h. 300 μL eluted phage was used to infect fresh XL1-Blue cells (Agilent 200228) at OD_600_ 0.6-0.8, and phage harvested from the infected cells. Positive selection with GFP with a C-terminal cyclic imide was followed by negative selection wherein the GFP-trap coupled to magnetic agarose (Bulldog Bio GTA020) was preincubated for 1 hour with GFP-His_6_, and the supernatant from the binding used to infect fresh XL1-Blue cells. Two more rounds of positive selection were performed, and plasmids were miniprepped from the infected outgrowth pellet with ZymoPURE Plasmid Miniprep Kit (Zymo Research D4212) according to kit instructions. PCR was performed on eluted plasmids by combining 12.5 ng plasmid, 1.25 μM of both forward and reverse primers (primers 95 and 96), and 1x Q5 High-Fidelity 2x Master Mix (New England Biolabs M0492) at 98°C for 30 seconds followed by 25 cycles of: 98°C for 10 seconds, 56.9°C for 30 seconds, and 72°C for 20 seconds, then 72°C for 2 minutes and a 4°C hold. PCR products were cleaned and Illumina Adaptors added as in 16s metagenomic library prep guide. Samples were sequenced using a MiSeq Nano flow cell with 250 bp-paired end reads and analyzed with Geneious Prime. In short, paired-end sequences were mapped to the reference sequence and variant frequency was calculated.

### Affinity maturation of CRBNmidi

Random mutagenesis was performed on target plasmids using flanking primers 97 and 98 with SfiI restriction sites inserted using Agilent GeneMorph II kit (Agilent Technologies 200550). The randomly mutagenized insert and pCOMB3xSS vector were digested by SfiI and the backbone gel extracted using the Monarch gel extraction kit (New England Biolabs T1020S). The backbone and insert were ligated at a 1:3 ratio using T4 DNA ligase (New England Biolabs M202T) at room temperature for 2 h. The ligation mix was transformed in 5-alpha (New England Biolabs C2987H) and random mutagenesis was verified by Sanger sequencing of the insert region of individual colonies. Verified libraries were electroporated into XL1-Blue cells (Agilent 200228) at 1700V using an Eppendorf electroporator. Cells were allowed to grow for 30 minutes, at which point 30 μL helper phage (M13KO7 New England Biolabs #N0315) were added and the cells allowed to grow for 1 hour. Additional media, kanamycin (50 μg/mL), and carbenicillin (100 μg/mL) were added and the cells allowed to grow overnight. Cells were pelleted at 4,000xg for 15 minutes at 4°C. Spent media was combined 5:1 with 2.5 M NaCl, 20% (w/v) PEG-8000 and incubated on ice for 20 minutes. Phage were pelleted at 12,000xg, 4°C, 30 minutes, and resuspended in PBS pH 7.4, 0.05% (v/v) Tween-20, 0.05% (w/v) bovine serum albimum. Resuspended phage was added to GFP-trap coupled to magnetic agarose (Bulldog Bio GTA020) preincubated for 1 hour with GFP-PFQYKcN. Phage and beads were incubated for 1 hour at 4°C. Phage were washed with 5% BSA in PBS or 5% BSA, 1% Triton-X in PBS by vortexing at lowest speed for 60 seconds and eluted by incubation with 0.5 M glycine-HCl pH 2.2 for 10 minutes at room temperature followed by neutralization with 0.15 volumes of 1 M Tris pH 9.1. 300 μL eluted phage was used to infect fresh XL1-Blue cells (Agilent 200228) at OD_600_ 0.6-0.8, and phage harvested from the infected cells after overnight incubation. Resuspended phage was added to GFP-trap coupled to magnetic agarose (Bulldog Bio GTA020) preincubated for 1 hour with GFP-PFQYKN. Phage were incubated with end-over-end motion at 4°C for 1 hour. Supernatant was transferred to GFP-trap coupled to magnetic agarose (Bulldog Bio GTA020) preincubated for 2 h with GFP-PFQYKcN. Phage and beads were incubated for 1 hour at 4°C. Phage were washed with specified washing buffer by vortexing on lowest speed for 90 seconds and eluted by incubation with 0.5 M glycine-HCl pH 2.2 for 10 minutes at room temperature followed by neutralization with 0.15 volumes of 1 M Tris pH 9.1. 300 μL eluted phage was used to infect fresh XL1-Blue cells (Agilent 200228) at OD_600_ 0.6-0.8, and phage harvested from the infected cells. This cycle repeated two additional times for a total of four rounds of selection. Plasmids were miniprepped from the infected outgrowth pellet with ZymoPURE Plasmid Miniprep Kit (Zymo Research D4212) according to kit instructions and whole plasmids were sequenced by Nanopore. Jupyter notebook was used to analyze data with coding assisted by ChatGPT (Supplementary Information).

### Virtual Screen of CRBNmidi linker

Simultaneous mutations of three residues within the GSG linker were performed on the lenalidomide-bound structure of CRBNmidi (PDB 8RQA)^23^ using Rosetta_Cartesian_ddG^47,48^ to screen 8000 possible variants. While Rosetta-based protocols are traditionally optimized for single-site mutations, the structural flexibility of the linker makes it a suitable region for multi-residue mutations without causing major disruptions.^49^ Among all tested variants, the GSG → RYD mutation resulted in the greatest reduction in free energy (ΔΔG = -9.6 kcal/mol).

### Overexpression and purification of GFP and eSrtA

Overexpression and purification of GFP and eSrtA were performed as described in Ichikawa et al.^12^

### eSrtA reactions

eSrtA reactions were performed as described in Ichikawa et al.^12^

### Overexpression and purification of TBD and mutations

All proteins were expressed in BL21(DE3) *E. coli* cells grown in LB supplemented with 50 μg/mL kanamycin. Protein expression was induced at OD_600_ 0.6-0.8 with 1 mM IPTG at 25°C for 4 h. After harvesting, the cells were suspended in PBS pH 7.4, 1% (v/v) Trition-X, cOmplete Protease Inhibitor (Roche) and lysed via sonication 20 seconds on, 10 seconds off at 27% amplitude on ice. The lysate was clarified by centrifugation at 21,130xg for 10 minutes at 4°C and loaded onto a HiTrap HP (Cytiva) column with NiSO_4_ bound. Elution occurred over a gradient between 25 mM and 500 mM imidazole in PBS. Fractions with protein were selected according to the A280 trace and concentrated to under 1 mL prior to size exclusion purification in tris-buffered saline using a pre-equilibrated S75 10/300 GL column.

### Overexpression and purification of CRBNmidi-derived mutations

Overexpression was performed as in Kroupova et al.,^23^ and purification protocol was adapted slightly. Harvested cells were frozen and resuspended in 20 mM HEPES pH 8.0, 500 mM NaCl, 50 μM ZnCl2, 0.5 mM TCEP, 0.05% Tween-20, 5 mM imidazole, 1 mM MgCl_2_, 10 μg/mL DNase, cOmplete Protease Inhibitor (Roche). Resuspended cells were incubated on ice for 30 minutes prior to lysis via sonication 20 seconds on, 10 seconds off at 27% amplitude on ice. The lysate was clarified by centrifugation at 21,130xg for 10 minutes at 4°C and loaded onto a HiTrap HP (Cytiva) column with NiSO_4_ bound. The column was washed with 15 CV 10 mM imidazole in 20 mM HEPES pH 8.0, 500 mM NaCl, 0.5 mM TCEP, followed by a gradient over 5 CV to 60 mM imidazole in 20 mM HEPES pH 8.0, 500 mM NaCl, 0.5 mM TCEP where it was held for 5 CV, then a gradient to 250 mM imidazole in 20 mM HEPES pH 8.0, 500 mM NaCl, 0.5 mM TCEP over 20 CV. Protein eluted around 100-150 mM imidazole and was pooled, aliquoted, and stored in elution buffer.

### Cloning, overexpression, and purification of UROD and UROD N367A

A geneblock coding for UROD was optimized for *E. coli* overexpression using IDT’s tool and purchased from Azenza. PCR was performed on a pET28a backbone (primers 83 and 84) and restriction digestion using DpnI (New England Biolabs R0176S), Xbal (New England Biolabs R0145S), and BamHI-HF (New England Biolabs R3136S) followed by DNA clean and concentrate (New England Biolabs T1130S). 50 ng backbone and 37.5 ng insert were ligated with T7 DNA ligase (New England Biolabs M0318), and the ligation reaction mixture transformed into 5-alpha (New England Biolabs C2987H). Single colonies were selected and sequences verified by Nanopore whole plasmid sequencing (Quintara Biosciences). Protein expression was adapted from Phillips et al.^36^ and Laterriere et al.^31^ The pET28a-UROD was expressed in BL21(DE3) *E. coli* cells grown in LB supplemented with 50 μg/mL kanamycin. Protein expression was induced at OD_600_ 0.5 with 500 μM IPTG at 37°C for 5 h. After harvesting, the cells were suspended in 300 mM NaCl, 50 mM NaPO_4_ pH 6.8, 10% glycerol, Trition-X, cOmplete Protease Inhibitor (Roche) and lysed via sonication 20 seconds on, 10 seconds off at 27% amplitude on ice. The lysate was clarified by centrifugation at 21,130xg for 10 minutes at 4°C and loaded onto a HiTrap HP (Cytiva) column with NiSO_4_ bound. Protein was washed with 20 CV 300 mM NaCl, 50 mM NaPO_4_ pH 6.8, 0.5 mM TCEP, 10% glycerol, followed by a linear gradient over 5 CV to 300 mM NaCl, 50 mM NaPO_4_ pH 6.8, 0.5 mM TCEP, 10% glycerol, 70 mM imidazole where a 25 CV wash occurred. Protein was eluted over a 40 CV gradient to 300 mM NaCl, 50 mM NaPO_4_ pH 6.8, 0.5 mM TCEP, 10% glycerol, 250 mM imidazole. Fractions with protein were selected according to the A280 trace and concentrated to under 1 mL prior to size exclusion purification in 50 mM Tris pH 7.5, 10% glycerol, 0.5 mM TCEP using a pre-equilibrated S75 10/300 GL column.

### Cloning, overexpression, and purification of GFP-UROD

pET28a-GFP was produced from pET28a-GFP-LPETG-His_6_ by PCR using the Q5 Site-Directed Mutagenesis Kit (New England Biolabs E0552S) to PCR amplify all parts of the plasmid except the LPETG-His_6_ sequence, digest the template, and ligate the PCR product (primers 79 and 80). The reaction mixture was transformed in Mach1 *E. coli*, grown overnight, miniprepped, and verified by Sanger sequencing. PCR to insert the His_6_ tag at the N-terminus was performed on the verified plasmid using the Q5 Site-Directed Mutagenesis Kit and primers 81 and 82 (New England Biolabs E0552S). The reaction mixture was transformed in Mach1 *E. coli*, grown overnight, miniprepped, and verified by Sanger sequencing. PCR was performed to cut the pET28a-GFP backbone to two pieces (primers 89-92), which were gel extracted using Monarch gel extraction kit (New England BIolabs T1010G). Assembly was performed by incubating the two backbone fragments with UROD amplified by PCR using primers 87 and 88 with HiFi DNA Assembly Master Mix (New England Biolabs E2621S) for 20 minutes at 50°C prior to transformation in 5-alpha (New England Biolabs C2987H) and verification by whole plasmid sequencing. The pET28a-GFP-UROD was expressed in BL21(DE3) *E. coli* cells grown in LB supplemented with 50 μg/mL kanamycin. Protein expression was induced at OD_600_ 0.5 with 500 μM IPTG at 20°C for 4.5 h. After harvesting, the cells were suspended in 300 mM NaCl, 50 mM NaPO_4_ pH 6.8, 10% glycerol, Trition-X, cOmplete Protease Inhibitor (Roche) and lysed via sonication 20 seconds on, 10 seconds off at 27% amplitude on ice. The lysate was clarified by centrifugation at 21,130xg for 10 minutes at 4°C and loaded onto a HiTrap HP (Cytiva) column with NiSO_4_ bound. Protein was washed with 20 CV 300 mM NaCl, 50 mM NaPO_4_ pH 6.8, 0.5 mM TCEP, 10% glycerol, followed by a linear gradient over 5 CV to 300 mM NaCl, 50 mM NaPO_4_ pH 6.8, 0.5 mM TCEP, 10% glycerol, 70 mM imidazole where a 25 CV wash occurred. Protein was eluted over a 40 CV gradient to 300 mM NaCl, 50 mM NaPO_4_ pH 6.8, 0.5 mM TCEP, 10% glycerol, 250 mM imidazole. Fractions with protein were selected according to the A280 trace and concentrated to under 1 mL prior to buffer exchange using a Zeba Spin column to 50 mM Tris pH 7.5, 10% glycerol, 0.5 mM TCEP.

### Electroporation of GFP-UROD

HEK293T cells were grown to 80-90% confluency in DMEM +/-prior to electroporation. Cells were detatched by trypsinization and washed with PBS prior to resuspension in PBS and counting. For each sample, an equal number of cells (7×10^5^) were aliquoted into a 1.7 mL tube and pelleted.

Electroporation mixes were prepared for each sample type by combining 41.6 μL Neon buffer R containing DMSO or 100 μM lenalidomide, 4.2 μL 50 μM mCherry protein as an internal standard, and 4.2 μL of 50 μM GFP-UROD with or without 17.5 hour preincubation with 150 μM PCMT1. A mock electroporation mix was also prepared using PBS in place of the GFP and mCherry proteins. Immediately prior to each electroporation, the PBS was removed and pelleted cells were resuspended in 10-12 μL of the electroporation mix. The sample was taken up in a 10 μL tip attached to a Neon pipette, and the pipette tip was submerged in a Neon cuvette containing 3 mL Neon buffer E. The sample was electroporated at 1150V, 20 msec, 2 pulses. The cells were then dispensed in 100 μL warmed PBS and flicked to mix. This process was repeated for each sample, with each tip used for 3 electroporation cycles. Cells were allowed to recover in PBS for at least 5 minutes after electroporating prior to centrifugation at 1,000xg for 5 min. The supernatant was removed, and cells were resuspended by pipetting in 50 μL colorless trypsin-EDTA. Resuspended cells were incubated at 37°C for 5 minutes. Trypsinization was quenched with the addition of 500 μL colorless DMEM +/+ containing DMSO or 100 μM lenalidomide and the mixture was transferred to a 24-well plate. Samples were incubated at 37°C, 5% CO_2_ for 6 h for preincubated samples or 42 h for samples not preincubated with PCMT1 prior to resuspension and analysis by flow cytometry (PE Texas Red and FITC on FACSymphony A3 Lite). To calculated normalized the normalized GFP level for WT HEK293T cells, the arithmetic mean of GFP and mCherry fluorescence intensities for each sample were corrected by subtracting the corresponding intensity values in the mock sample; then, these corrected values were used to calculate a GFP/mCherry fluorescence intensity ratio for each sample. For HEK293T CRBN KO cells, the arithmetic mean of the GFP fluorescence intensity for each sample was corrected by subtracting the corresponding intensity value in the mock sample. For both cell types, the resulting values were normalized to the mean GFP/mCherry or GFP value for that GFP-UROD in the presence of lenalidomide.

### Site directed mutagenesis for point mutations

Site directed mutagenesis was performed using QuikChange Lightening Site-Directed Mutagenesis Kit (Agilent Technologies 210518 or 210519) using primers designed with Agilent’s Primer Design Program (primers 1-78. 85-86, and 93-94).

### Western Blotting

Samples were mixed with 5x SDS-PAGE loading buffer (1x concentration: 50 mM Tris–HCl, 2% SDS, 10% glycerol, 1% β-mercaptoethanol, 0.02% bromophenol blue) and boiled at 95°C for 5 minutes. The denatured samples (8–10 µL) were loaded on a 12% Criterion TGX 26-well precast gel and run at 180 V, 400 mA in Tris/Glycine/SDS buffer. Gels were dry transferred using iBlot 2 or powerblotter at 25 V for 7 minutes. Total protein was imaged by Ponceau staining. Membranes were blocked in 5% (w/v) bovine serum albimum or nonfat dry milk powder before addition of primary antibody or cerebody. Cerebody blots were performed with a final concentration of 5 μM protein in 5% (w/v) nonfat dry milk in TBST unless specified. All primary antibodies were incubated overnight. Membranes were washed 3x, 5 minutes in TBST, incubated with secondary antibody in blocking buffer 1 hour at 22°C with motion, then washed 3x, 5 min prior to visualization by IR800 or chemi channel on an Azure imager.

### Overexpression and purification of GFP-GLUL, GLUL and GLUL N373A

Overexpression of GLUL and GLUL N373A were performed as described in Zhao et al.^15^ BL21(DE3) cells were transformed with the validated plasmid and used to inoculate overnight cultures of LB + 100 µg/mL carbenicillin. Large-scale overexpression cultures (1 L×5) were inoculated with overnight culture diluted 1:100. The overexpression cultures were incubated at 37°C with shaking at 220 rpm until the OD600 was approximately 0.6, at which point the temperature was reduced to 16 °C and IPTG was added to a final concentration of 0.42 mM. The cultures were incubated for approximately 24 h prior to collecting the cells by centrifugation. If desired, the cell pellets were flash frozen and stored at –80°C.

Cell pellets were resuspended in 8.3 mL of 20 mM imidazole, 1× protease inhibitor, 1% Triton-X 100/PBS per pellet. Lysates were sonicated (6 sec on, 60 sec off, 5 min total, 70% amplitude using a Fisherbrand model 120 sonicator) on ice and clarified by centrifugation (20,000 × g, 4 °C, 10 min) and syringe filtration (0.45 µm). The His-tagged protein was first purified on a Ni-bound 1 mL HiTrap Chelating HP column, equilibrating and washing with 20 mM imidazole/PBS and eluting with a gradient to 500 mM imidazole/PBS (both pH 8.0). Protein-containing fractions were concentrated to < 1 mL and further purified on a S200 10/300 GL column, pre-equilibrated and run with PBS (pH 7.4). Protein-containing fractions were collected and concentrated with a 10 kDa MWCO spin filter, and the concentration determined by A280 (ε = 61,475 M^-1^ cm^-1^). The fractions with the most prominent peak shape were aliquoted, flash frozen and stored at –80 °C.

### Overexpression and purification of PCMT1 and PCMT1 S60A

Overexpression of PCMT1 and PCMT1 S60A were performed as described in Zhao et al.^15^ BL21(DE3) cells were transformed with the validated plasmid and used to inoculate overnight cultures of LB + 50 µg/mL kanamycin. Large-scale overexpression cultures (750 mL×4) were inoculated with overnight culture diluted 1:100. The overexpression cultures were incubated at 37°C with shaking at 200 rpm until the OD600 was approximately 0.6, at which point the temperature was reduced to 16 °C and IPTG was added to a final concentration of 0.4 mM. The cultures were incubated for approximately 22 h prior to collecting the cells by centrifugation. If desired, the cell pellets were flash frozen and stored at –80°C.

Cell pellets were resuspended in 8.3 mL of 20 mM imidazole, 1× protease inhibitor, 1% Triton-X 100/PBS per pellet. Lysates were sonicated (30 sec on, 10 sec off, 5 min total, 25% amplitude using Branson model 250 sonicator) on ice and clarified by centrifugation (20,000 × g, 4 °C, 10 min) and syringe filtration (0.45 µm). The His-tagged protein was first purified on a Ni-bound 1 mL HiTrap Chelating HP column, equilibrating and washing with 20 mM imidazole/PBS and eluting with a gradient to 500 mM imidazole/PBS. Protein-containing fractions were concentrated to < 1 mL and further purified on a S75 10/300 GL column, pre-equilibrated and run with TBS. The fractions from the second apparent peak, validated to have the best enzymatic activity, were collected and concentrated with a 10 kDa MWCO spin filter, and the concentration determined by A280 (ε = 21,555 M^-1^ cm^-1^). The protein was aliquoted, flash frozen, and stored at –80 °C.

### PCMT1 Incubations

PCMT1 reaction conditions were adapted from in Zhao et al.^15^ In brief, 200 μM SAM, 10 μM PCMT1, and 10–50 μM purified protein or 1-2 mg/mL lysate were incubated at 37°C for 12-14 h prior to quenching via addition of 5x BME gel loading buffer and heating at 95°C for 5 minutes or proceeding directly to precipitation for GLUL, or proceeding directly to TR-FRET for GFP-GLUL. For UROD, 200 μM SAM, 50 μM UROD, and 150 μM PCMT1 were incubated at 37°C for 17.5 h prior to TR-FRET, Western blotting, or digestion unless otherwise specified.

### Cell culture

Human-derived cell lines were cultured in DMEM supplemented with 10% heat-inactivated fetal bovine serum (FBS) and 1xpenicillin-streptomycin. Cells were grown at 37°C in a humidified atmosphere with 5% CO_2_. For the collection of cell pellets, cells were dissociated with trypsin and collected by centrifugation at 500xg, 22°C, 3 min, followed by a PBS wash. Pellets were used immediately or stored at -80°C until use. Cultured cell lysates were prepared by resuspending cell pellets Pierce IP Lysis Buffer (Thermo Scientific 87787) supplemented with 1x protease/phosphatase inhibitor cocktail (CST 5872) by pipetting. Resuspended cells were incubated on ice for 10 minutes. The lysate was clarified by centrifugation at 21,130xg for 10 minutes at 4°C, and protein concentration measured by BCA.

### Mouse sample preparation

C57BL/6J female mice were obtained from The Jackson Laboratory. Mice were maintained in an Association for Assessment and Accreditation of Laboratory Animal Care (AAALAC) approved animal facility at Harvard University. Procedures were approved by the Institutional Animal Care and Use Committee of all institutions. Normal brain, kidney, liver, spleen, and hearts were collected from mice aged 12.5 weeks. Dissected tissues were flash frozen and stored at -80°C until use. Portions of tissue samples were placed in T-PER (Thermo Scientific 78510) supplemented with 1x protease/phosphatase inhibitor cocktail (CST 5872) and sonicated at 12% amplitude until mixture appeared homogenous. The lysate was clarified by centrifugation at 21,130xg for 10 minutes at 4°C, and protein concentration measured by BCA.

### Affinity enrichment with cerebody

If applicable, compound or an equivalent volume of DMSO was added to samples for precipitation, followed by addition of cerebody to a final concentration of 5 μM in an initial 1-2 mg/mL sample, not including exogenous PCMT1 if applicable. Washed Sigma Anti-FLAG^®^ M2 magnetic beads (M8823) were added at 4.5 μL packed beads/100 μL initial sample. Samples with cerebody and beads were incubated on a roller at room temperature for 80-90 minutes. Samples were briefly centrifuged, and beads captured using a magnetic rack. Supernatant was removed and 80 μL 1% (v/v) Trition X in PBS was added per 100 μL initial sample. Beads were vortexed at the lowest speed for 60 seconds. Samples were briefly centrifuged, and beads captured using a magnetic rack and the supernatant removed. 1x gel loading buffer or 5% SDS, 50 mM TEAB pH 7.4 at around 20 μL/100 μL initial sample was added, and samples heated at 95°C for 5-10 minutes. Samples were centrifuged and the magnetic beads captured. Supernatant was transferred to a clean tube for Western blotting or further analysis.

### Mass spectrometry of cerebody affinity enrichments

Samples eluted in 5% SDS, 50 mM TEAB pH 7.4 were reduced by addition of 20 mM dithiothreitol at 22°C for 30 minutes then alkylated by addition of 40 mM iodoacetamide and incubation in the dark at 22°C for 30 minutes. The reaction was quench by addition of phosphoric acid to 1.2% final concentration by volume. 900 μL S-trap buffer (90% methanol, 100 mM TEAB pH 7.4) was added to each sample prior to transfer to an S-trap micro column. Using a vacuum manifold, the columns were washed 3 times with 150 μL S-trap buffer. 1 μg of trypsin resuspended in 25 μL 50 mM TEAB pH 7.4 was added to each column and incubated at 47°C for 2 h. The digested peptides were eluted by sequential addition of 40 μL 50 mM TEAB pH 7.4, 40 μL 0.2% (v/v) formic acid, and 40μL 0.2% formic acid, 50% acetonitrile (v/v) with each elution being collected by centrifugation (4000xg, 22°C, 1 min) to a clean tube. The eluted samples were concentrated to dryness in a vacufuge and resuspended in 25 μL ddH2O. To each sample, 10 μL TMT reagent was added and incubated at 22°C for 1 hour. Tris pH 7.6 was added to a final concentration of 167 μM and incubated for 15 minutes at 22°C to quench the TMT reagents. The TMT-labeled samples were combined and dried in a vacufuge. The dried sample was resuspended in 300 μL 0.1% trifluoroacetic acid (v/v) in water and fractionated by the Pierce High pH reversed phase peptide fractionation kit (84868). The peptides were eluted sequentially by ddH2O, 2x 5% acetonitrile/0.1% TEA, 10% acetonitrile/0.1% TEA, 15% acetonitrile/0.1% TEA, 25% acetonitrile/0.1% TEA, 50% acetonitrile/0.1% TEA, and acidified to a final concentration of 0.164% formic acid. The water and 5% acetonitrile/0.1% TEA fractions were excluded from LC-MS/MS analysis. The other fractions were concentrated to dryness by vacufuge and resuspended in 30μL 0.1% formic acid prior to LC-MS/MS analysis. The LC-MS/MS experiment was performed on an Orbitrap Eclipse Tribrid Mass Spectrometer coupled with a Vanquish Neo HPLC system (Thermo scientific) at the Harvard Center for Mass Spectrometry. The TMT labelled peptides were first trapped on a trapping cartridge (300µm x 5mm PepMap™ Neo C18 Trap Cartridge, Thermo scientific) prior to separation on a silica-chip-based micropillar column (µPAC, C18 pillar surface, 50 cm bed, Thermo scientific). The column oven temperature was maintained at 35 °C. Peptides were eluted using a multistep gradient at a flow rate of 0.3 µL/min over 180 min. The Orbitrap Eclipse MS was operated in data dependent acquisition (DDA) mode with positive polarity in combination with high field asymmetric waveform ion mobility spectrometry (FAIMS) pro interface. FAIMS compensation voltages (CV) of -40V, -55V, -70V were combined in a single run with a cycle time of 1.5 sec. FAIMS carrier gas flow rate was at 4.6L/min. Spray voltage was set to 2.1 kV and heated capillary temperature at 275 °C. Full scan was performed in the range of 400-1,600 m/z at a resolution of 120,000, radio frequency (RF) lens 30%, normalized automatic gain control (AGC) target at standard and maximum injection time set to auto. Precursors were isolated in a window of 0.7 m/z and charge states from two to six. Fragmentation was performed by higher-energy collisional dissociation (HCD) using normalized collision energy (NCE) of 38% at the resolution of 50,000. The normalized AGC target was set to 250%, and a maximum ion injection time set to 200 ms. Dynamic exclusion was enabled with a mass tolerance of 10 ppm and exclusion duration of 60s.

Searches were performed with the following guidelines: semitrypsin digestion at the peptide N-terminus only to search for cleavage peptides and full trypsin digestion to search for deamidated peptides and quantify total protein level; fewer than two missed cleavages; static carboxyamidomethylation of cysteine residues (+57.021 Da); static TMTpro 16-plex labeling (+304.207 Da) at lysine residues and N termini; variable oxidation on methionine residues (+15.995 Da); variable dehydration on asparagine and glutamine residues at C-terminus only (– 18.015 Da). To search for deamidation sites, the variable dehydration modification was replaced with variable deamidation on asparagine residues (+0.984 Da). The TMT reporter ions were normalized to the total peptide amount. For the quantification of total proteins, the data were filtered to include only master proteins with high protein FDR confidence, more than one peptide, and more than one protein unique peptide, and exclude all contaminant proteins, with the cerebody quantification serving as a normalization factor. The abundance ratios and their associated *P*-values were calculated by one-way ANOVA with the Tukey HSD post hoc test.

### Mass spectrometry of UROD

25 μL 50 μM UROD incubated with or without PCMT1 for 17.5 h was buffer exchanged using a 0.5 mL Zeba spin column to 100 mM ammonium bicarbonate pH 7.4, and the volume was brought to 80 μL with a final concentration of 0.1% Rapigest. The samples were reduced by addition of dithiothreitol (10 μL of a 45 mM stock) for 30 minutes at 22°C then alkylated by addition of iodoacetamide (10 μL of a 150 mM stock) for 30 minutes at 22°C in the dark. The samples were digested with 1 μg of Glu-C for 2 h at 37°C, then acidified by addition of trifluoroacetic acid (5 μL of a 12% solution) to a final concentration of 0.5%. Samples were incubated at 37°C for 30 minutes prior to centrifugation at 13,000xg for 10 minutes. Supernatant was transferred to a clean tube and concentrated to dryness by vacufuge. Samples were desalted using the Pierce peptide desalting spin columns (ThermoFisher 89851), and peptides eluted in 0.1% formic acid/50% acetonitrile. They were concentrated to dryness by vacufuge and resuspended in 20 μL 0.1% formic acid prior to LC-MS/MS analysis. The same Orbitrap Eclipse with Vanquish Neo LC-MS/MS system as mentioned above was used for this analysis. The peptide samples were first trapped on a trapping cartridge prior to separation on a µPAC silica-chip-based micropillar column. The conditions of the trap and column were as described above. Peptides were eluted using a gradient at a flow rate of 300 nL/min over 90 min. The mobile phase A and weak wash liquid was water with 0.1% FA, and mobile phase B was acetonitrile with 0.1% FA, and strong wash liquid was 80% acetonitrile with 0.1% FA. The mobile phase gradient consisted of a linear 69-min gradient from 1% to 33% mobile phase B, followed by a 13-min increase to 46% B, a further 8-min plateau phase at 95% B. The autosampler temperature was 7 °C. The column “Fast equilibration” was enabled. The Orbitrap Eclipse MS was operated in positive electrospray ionization mode with a voltage of 2.1 kV and the capillary temperature at 275 °C. FAIMS compensation voltage was set at -45V with a cycle time of 3 sec. The target selected ion monitoring (tSIM) was performed at a resolution of 120,000, an isolation window of 1 m/z, a RF lens of 30%, with normalized AGC target 200% and maximum injection time 300ms. The target mass inclusion list is below.

| m/z | z | t start (min) | t stop (min) |
| --- | --- | --- | --- |
| 728.7348 | 3 | 0 | 90 |
| 546.8029 | 4 | 0 | 90 |
| 437.6438 | 5 | 0 | 90 |
| 722.7312 | 3 | 0 | 90 |
| 542.3002 | 4 | 0 | 90 |
| 434.0417 | 5 | 0 | 90 |
| 729.9237 | 2 | 0 | 90 |
| 486.9515 | 3 | 0 | 90 |
| 720.9184 | 2 | 0 | 90 |
| 480.948 | 3 | 0 | 90 |

Dynamic exclusion was enabled with a mass tolerance of 10 ppm and exclusion duration of 1s. Fragmentation of target precursors was performed by HCD using NCE of 30% at the resolution of 60,000. The normalized AGC target was set to 100%, and a maximum ion injection time set to 150 ms.

## Acknowledgements

We thank N. Vallavoju, A.K.D. Porter, Q. Zhu, and M. Huffman for helpful discussions and input; C. Hartmann from the Harvard University Bauer Core Facility; and the 2021 Antibody Engineering & Display Technologies Course at Cold Spring Harbor Laboratory for guidance with phage display. CRBN/DDB1 was a generous gift from Boehringer Ingelheim. Support from the NIH (R01 GM141406, C.M.W.; F31, AR083715, R.F.), the Starr Foundation (C.M.W.), the Mark Foundation for Cancer Research (C.M.W.), and New York Stem Cell Foundation (Y.-C.H.) is gratefully acknowledged. Research in the Ciulli laboratory on studying and targeting Cullin RING E3 ligases receives funding from the Cancer Grand Challenges partnership funded by Cancer Research UK, Institut National Du Cancer (INCa), and KiKa (Children Cancer Free Foundation); and the Innovative Medicines Initiative 2 (IMI2) Joint Undertaking under grant agreement No 875510 (EUbOPEN project to A.C.) from the European Union’s Horizon 2020 research and innovation programme. The IMI2 Joint Undertaking receives support from the European Union’s Horizon 2020 research and innovation program, European Federation of Pharmaceutical Industries and Associations (EFPIA) companies, and associated partners KTH, OICR, Diamond, and McGill.

## Author Contributions

H.C.L, Y.L., N.C.P., W.X., F.K., M.C., R.M., and C.M.W. designed research. H.C.L., Y.L., N.C.P., and M.C. performed research. H.C.L., Y.L., Z.Z., W.X., A.K., D.Z., T.L., R.F., E.Y.F., S.X., Y.-C.H., and A.C. contributed reagents or materials. H.C.L., Y.L., N.C.P., W.X., M.C., R.M., and C.M.W. analyzed data. H.C.L. and C.M.W. wrote the paper; all coauthors reviewed the paper.

## Declaration of Interest

R.M. and N.C.P. are inventors on patent applications related to the CoraFluor TR-FRET technology used in this work. The Woo Lab receives or has received sponsored research support from Amgen, Ono Pharmaceuticals, and Merck. The Ciulli laboratory receives or has received sponsored research support from Almirall, Amgen, Amphista Therapeutics, Boehringer Ingelheim, GlaxoSmithKline, Eisai, Merck KGaA, Nurix Therapeutics, Ono Pharmaceuticals, and Tocris-Biotechne. A.C. is a co-founder and shareholder of Amphista Therapeutics, a company that is developing targeted protein degradation therapeutic platforms.

## Data Availability

Proteomics data have been deposited to the PRIDE repository with the dataset identifier PXD061728. All other data are included in the main manuscript, SI Appendix, and Supplemental Tables 1-8.

---

**Extended Data Figure 1.**
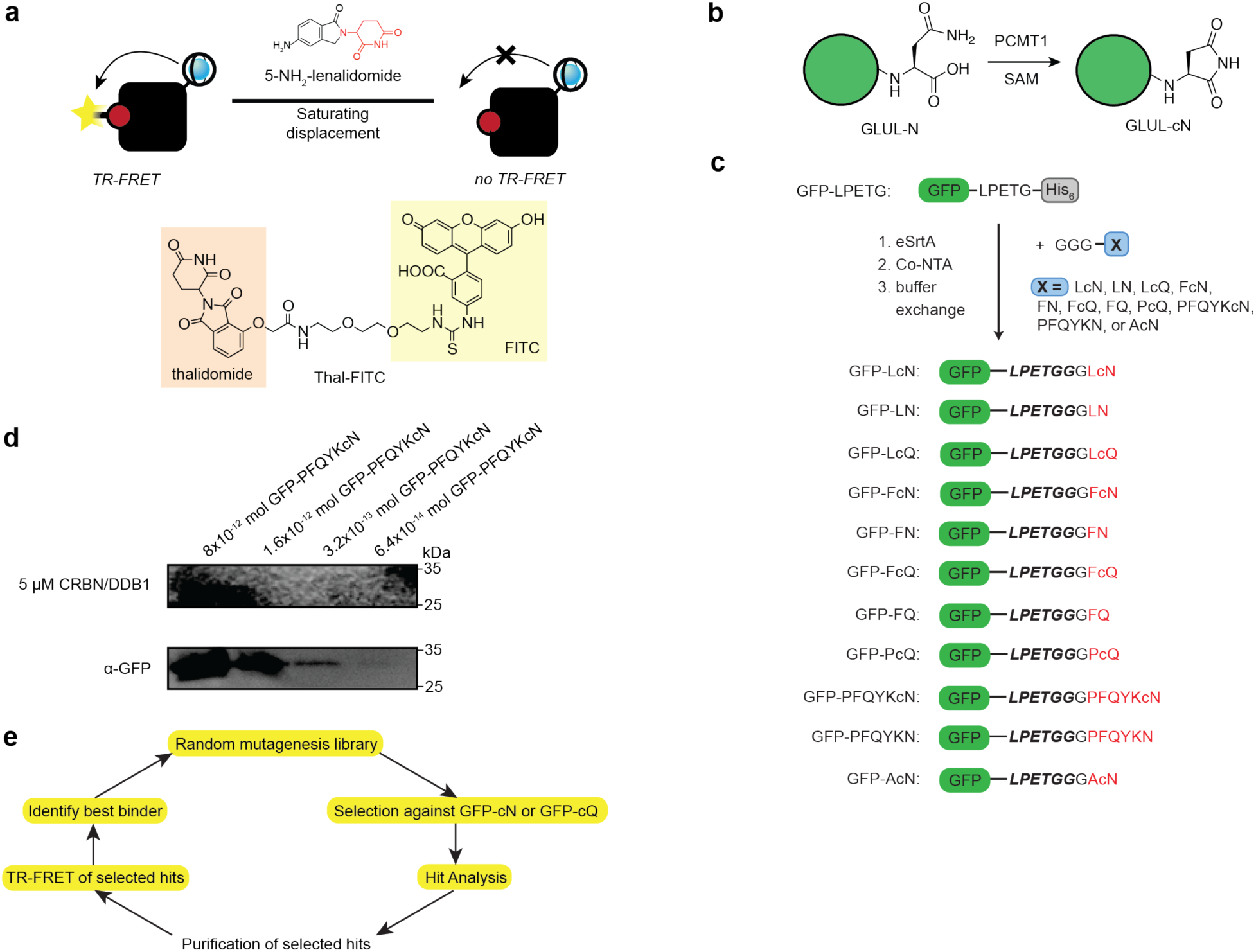
Phage display of TBD and application of mutations to CRBNmidi. **(a)** Illustration of association TR-FRET assay. **(b)** Overview of PCMT1-catalyzed C-terminal cyclic imide formation on GLUL. **(c)** Overview of eSrtA reaction to engineer semisynthetic GFPs used in this study. **(d)** Western blot of GFP-PFQYKcN using CRBN/DDB1 in place of a primary antibody and using α-GFP antibody. **(e)** Workflow of TBD evolution.

**Extended Data Figure 2.**
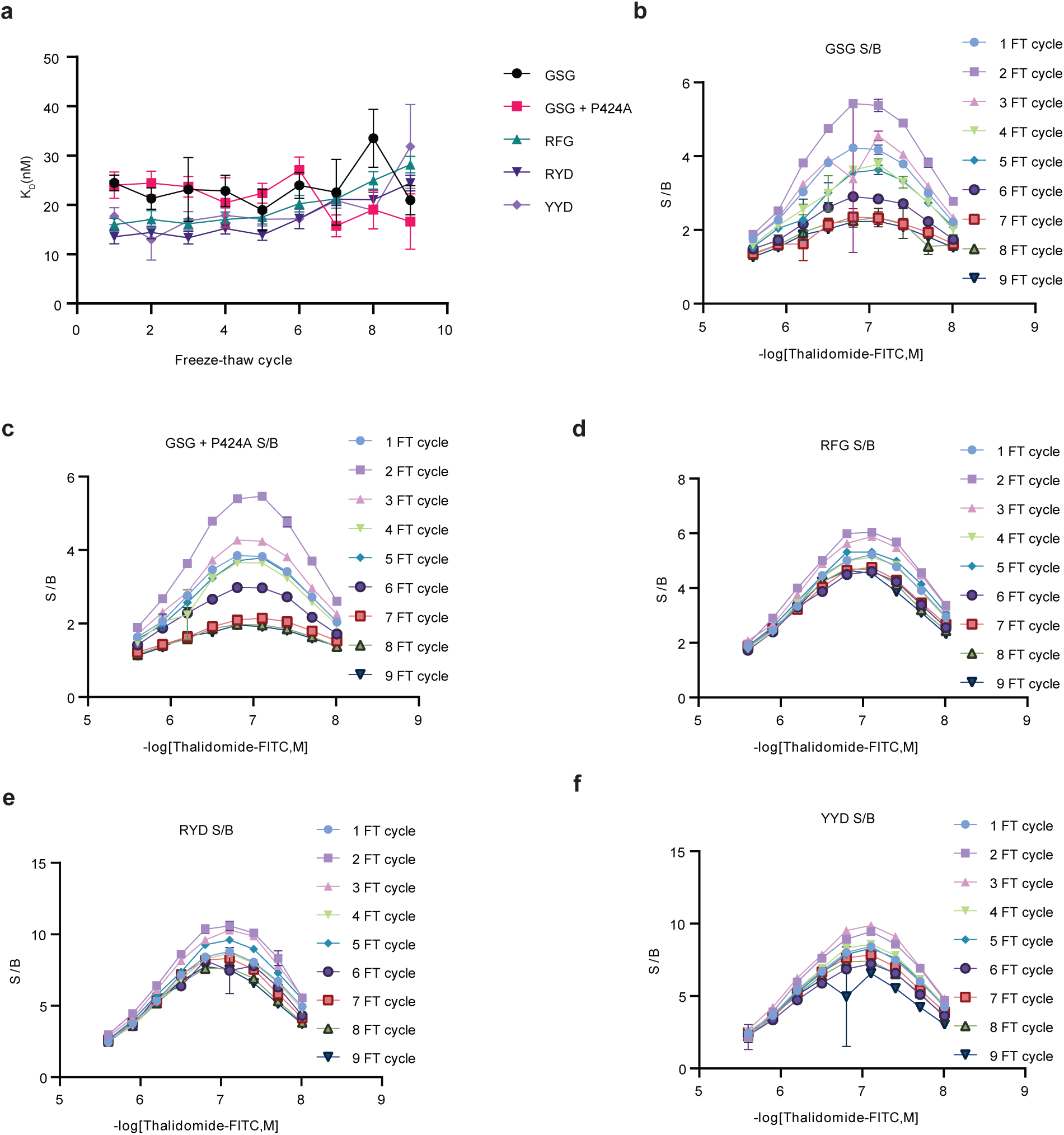
Optimization of cerebody linker from CRBNmidi. **(a)** K_D_s of CRBNmidi mutations based on linker screen over 9 freeze-thaw cycles. **(b)** Signal to background ratios of CRBN midi with the native GSG linker across 9 freeze thaw cycles and different thal-FITC concentrations. **(c)** Signal to background ratios of CRBN midi with the native GSG linker and the mutation corresponding to P424A across 9 freeze thaw cycles and different thal-FITC concentrations. **(d)** Signal to background ratios of CRBN midi with an RFG linker across 9 freeze thaw cycles and different thal-FITC concentrations. **(e)** Signal to background ratios of CRBN midi with a RYD linker across 9 freeze thaw cycles and different thal-FITC concentrations. **(f)** Signal to background ratios of CRBN midi with a YYD linker across 9 freeze thaw cycles and different thal-FITC concentrations.

**Extended Data Figure 3.**
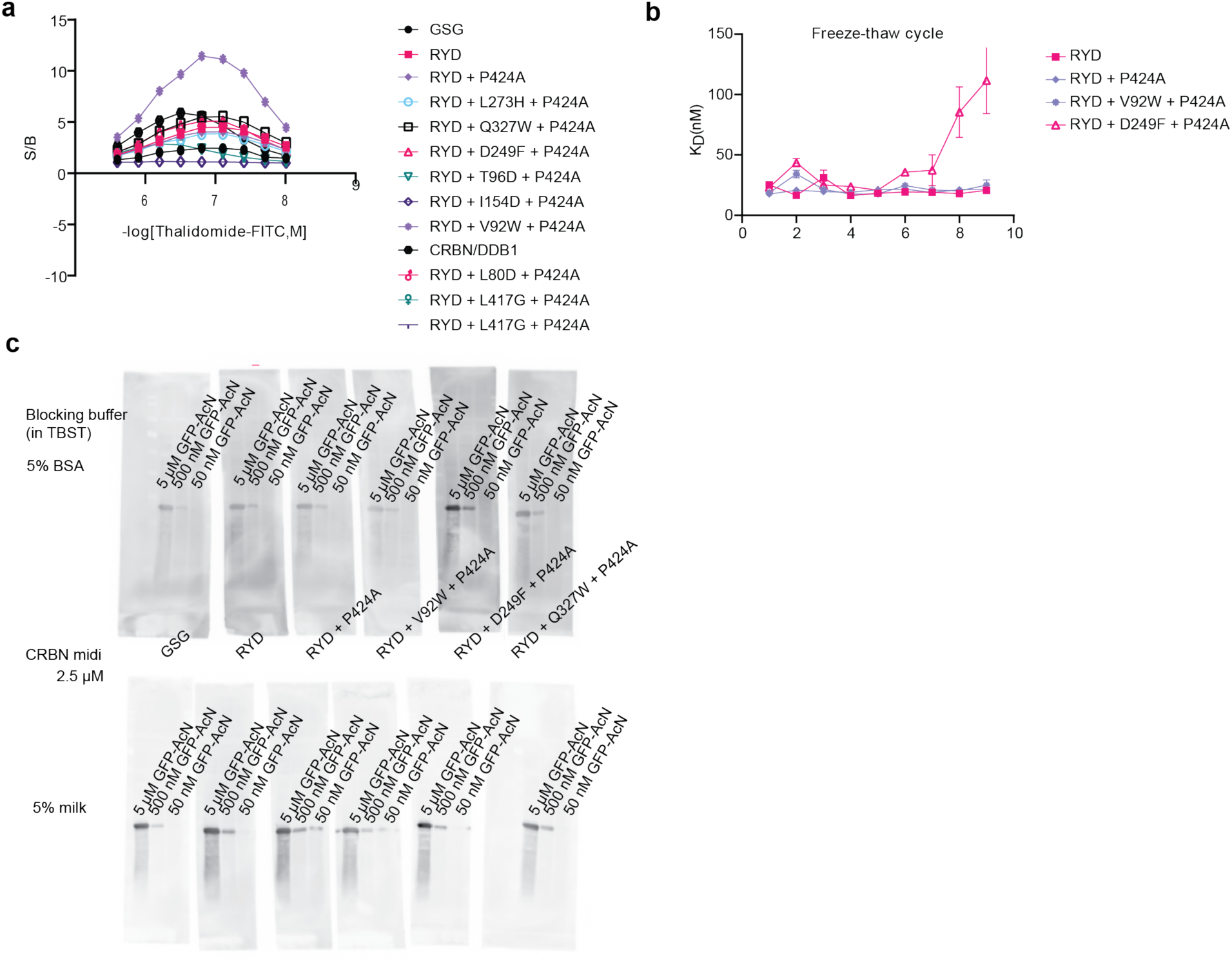
Optimization of cerebody from CRBNmidi-RYD. **(a)** Signal to background ratios of different CRBNmidi constructs. **(b)** Calculated K_D_ values of CRBN midi mutations based on phage display over 9 freeze thaw cycles. **(c)** Western blotting of different concentrations of GFP-AcN with CRBNmidi mutations in 5% milk or 5% BSA blocking buffer.

**Extended Data Figure 4.**
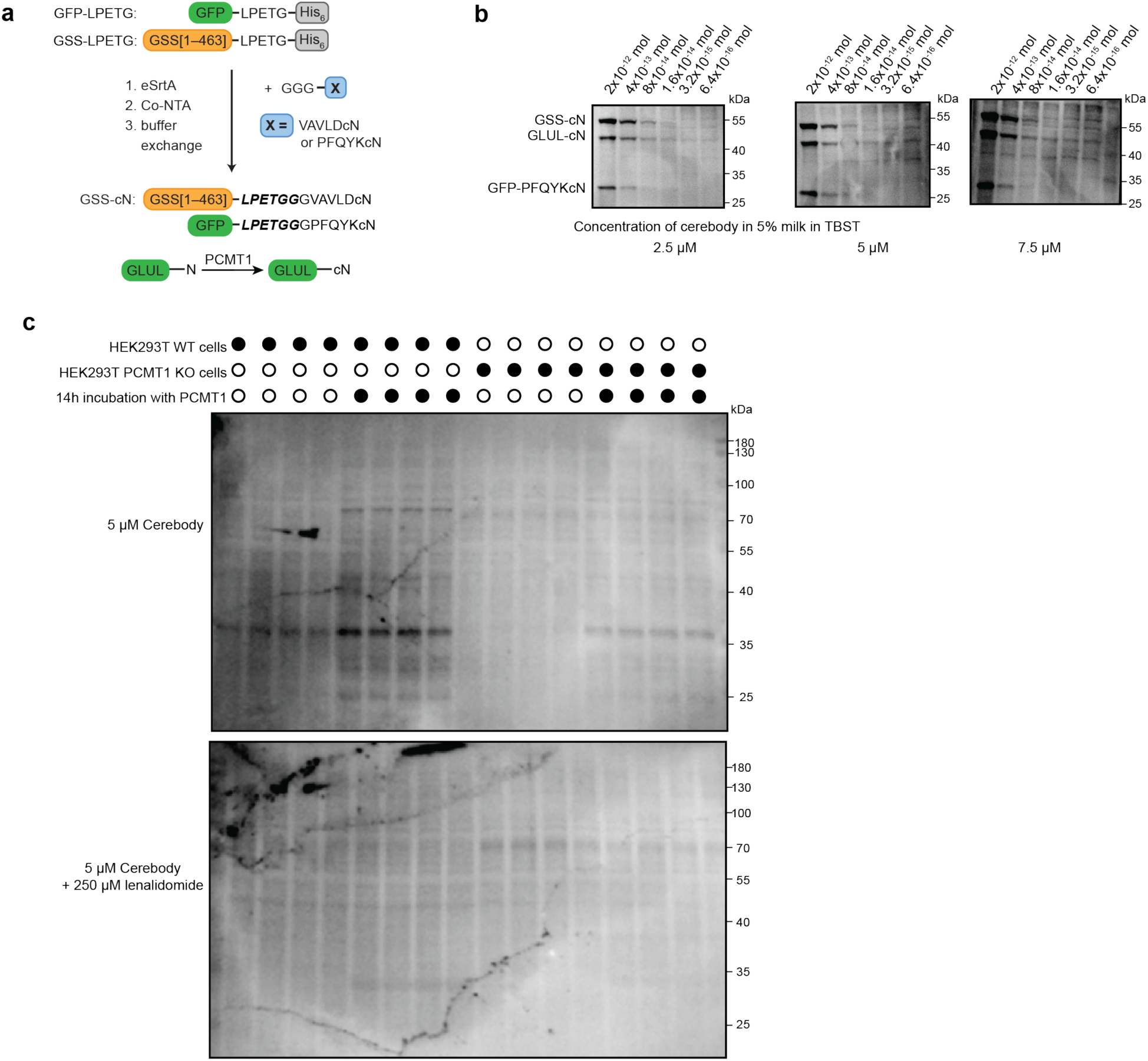
Western blot with cerebody. **(a)** Overview of C-terminal cyclic imide modified proteins. **(b)** Western blots of serial dilutions of GSS-cN, GFP-PFQYKcN, and GLUL incubated with PCMT1 for 14 h spiked in HEK293T lysate using different concentrations of cerebody in 5% milk in TBST as the primary antibody. **(c)** Cerebody blot of HEK293T WT and PCMT1 KO lysates with and without PCMT1 incubation.

**Extended Data Figure 5.**
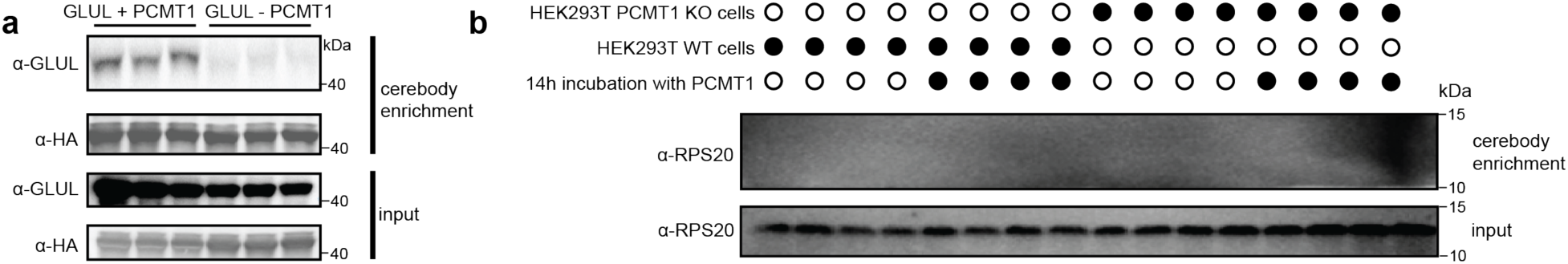
Cerebody enrichment. **(a)** Blot of GLUL with or without PCMT1 incubation that was spiked into HEK293T lysate prior to cerebody enrichment. **(b)** Blot of cerebody enrichment of HEK293T WT and PCMT1 KO cells with or without PCMT1 incubation for RPS20.

